# MISTR: A conserved <u>MI</u>tochondrial <u>ST</u>ress <u>R</u>esponse network revealed by signatures of evolutionary conflict

**DOI:** 10.1101/2020.01.25.919811

**Authors:** Mahsa Sorouri, Tyron Chang, Palmy Jesudhasan, Chelsea Pinkham, Nels C. Elde, Dustin C. Hancks

## Abstract

Host-pathogen conflicts leave genetic signatures of variation in homologous host genes. Using these “molecular scars” as a guide, we discovered a vertebrate-specific <u>MI</u>tochondrial <u>ST</u>ress <u>R</u>esponse circuit (MISTR). MISTR proteins are associated with electron transport chain factors and activated by stress signals such as interferon-gamma and hypoxia. Upon stress, ultraconserved miRNAs downregulate MISTR1 followed by replacement with paralogs MISTR <u>A</u>nti<u>V</u>iral (MISTRAV) or MISTR <u>H</u>ypoxia (MISTRH), depending on the insult. While cells lacking MISTR1 are more sensitive to apoptotic triggers, cells lacking MISTRAV or expressing the poxvirus-encoded vMISTRAV exhibit resistance to the same insults. Rapid evolution signatures across primate genomes for *MISTR1* and *MISTRAV* indicate ancient and ongoing conflicts with pathogens. MISTR proteins are also found in plants, yeasts, and an algal virus indicating ancient origins and suggesting diverse means of altering mitochondrial function under stress. The discovery of MISTR circuitry highlights the use of evolution-guided studies to reveal fundamental biological processes.

## INTRODUCTION

Innate immunity is a critical frontline host defense mechanism in response to pathogen infection. At the onset of infections in vertebrates, a set of more than 400 genes are transcriptionally upregulated by interferon and thus termed interferon-<u>s</u>timulated <u>g</u>enes (ISGs). ISGs display diverse functions such as activation of cell death programs and recruitment of immune cells (e.g. dendritic cells (Schneider et al., 2014; Schoggins, 2014). Although the identities of many of these genes are established, the functions of the majority of these gene products as well as their relationship with other cellular factors is unknown (Schoggins et al., 2011; 2014). To dissect the function of poorly characterized ISGs, we used a comparative approach to identify signatures of positive selection consistent with important functional roles in immune responses (Daugherty and Malik, 2012). Specifically, we sought to identify ISGs lacking known functions with hallmarks of repeated conflicts with pathogens, like those detected for interferon-inducible double-stranded RNA sensor <u>o</u>ligo<u>a</u>denylate <u>s</u>ynthetase <u>1</u> (OAS1), which in addition to signatures of rapid evolution (Hancks et al., 2015; Mozzi et al., 2015) are also encoded in virus genomes (Darby et al., 2014).

Viruses can encode proteins that mimic host proteins to manipulate cellular functions and inactivate immune defenses. This form of mimicry is commonly achieved by the acquisition of a host-coding sequence through horizontal gene transfer (HGT) followed by subfunctionalization via cycles of mutation and selection (Elde and Malik, 2009). Importantly, many viral mimics can be identified based on residual sequence identity (Spector et al., 1978). Along with inhibitors of immune function, mimics for master regulators of cellular functions such as *vSRC*, *vMYC*, and *vRAS* have been identified in virus genomes [reviewed in (Bishop, 1985)]. Our studies here stem from the identification of a viral ortholog for the ORFan ISG, *C15ORF48* [also known as *normal mucosal esophageal-specific gene product 1* (*NMES1*) (Zhou et al., 2002), mouse *AA467197*] - hereafter *<u>MI</u>tochondrial <u>ST</u>ress <u>R</u>esponse <u>A</u>nti<u>V</u>iral* (*MISTRAV*) - which is encoded by the large double-stranded DNA (dsDNA) virus squirrelpox, *<u>v</u>iral MISTRAV* (*SQPV078/vMISTRAV*).

Pathogen proteins mimicking host functions often directly bind host factors during infection to modulate cellular functions (Elde et al., 2008). Over evolutionary time, this action can impose strong selective pressure on the infected host, resulting in the increased frequency of host genetic variants in the population less susceptible to binding by a pathogen-encoded inhibitor (Daugherty and Malik, 2012). Recurrent rapid evolution resulting from genetic conflicts can be observed at the sequence level when the rate of nonsynonymous amino acid substitutions relative to the rate of synonymous substitutions (dN/dS) is greater than one when comparing orthologous proteins from closely related species.

Here, we focus on the rapidly evolving, interferon-gamma inducible MISTRAV which is encoded by a poxvirus. Our dissection of MISTRAV function unexpectedly revealed a stress-response circuit involving its paralogs *MISTR1* [also known as *NADH dehydrogenase ubiquinone 1 alpha subcomplex subunit 4* (*NDUFA4*)] and *MISTR <u>H</u>ypoxia* (*MISTRH*) [also known as *NADH dehydrogenase ubiquinone 1 alpha subcomplex subunit 4 like-2* (*NDUFA4L2*)], which are linked through regulation by the ultraconserved miRNAs *miR-147b* and *miR-210*. Localization analysis indicates MISTRAV and the virus-encoded vMISTRAV are mitochondrial proteins in agreement with paralogs MISTR1 (Balsa et al., 2012) and MISTRH (Tello et al., 2011) being putative supernumerary electron transport chain (ETC) factors.

Functional analysis in cell lines shows that loss of MISTRAV is associated with a reduction in apoptosis – a fundamental host defense to block pathogen replication. Correspondingly, a mutation resulting in a > 30-fold increase in levels of the *MISTRAV*-embedded *miR-147b* triggers a more robust activation of apoptosis in response to the cell death agonist staurosporine. Genetic and functional analysis reveal that the rapidly evolving paralog of *MISTRAV* - *MISTR1* - is a major target of the ultraconserved *miR-147b* as well as the hypoxia-inducible *miR-210* (Huang et al., 2009) which targets the same microRNA response element as *miR-147b*. Furthermore, we show that vMISTRAV can counteract triggers of apoptosis, consistent with the ability of viruses to counteract host defenses mediated by MISTR.

We propose a model for the vertebrate-specific <u>MI</u>tochondrial <u>ST</u>ress <u>R</u>esponse circuit (MISTR). *MISTR* genes exhibit a wide distribution of variation including homologs in plants, animals, and parasites, repeated duplications of paralogs between species, and a MISTR homolog in a giant DNA virus that infects algae. In addition to augmenting host immune defenses, MISTR may be a modular system with the capacity to respond to diverse stressors through regulation by specific miRNAs that downregulate MISTR1, while concurrent induction of MISTR paralogs replace MISTR1 to shape the mitochondrial response to perturbations.

## RESULTS

### *MISTR* proteins are encoded by highly diverged large DNA viruses

Human *MISTRAV* (*C15ORF48, NMES1*) is an eighty-three amino acid (AA) protein with a short N-terminus and a longer C-terminus demarcated by an intervening predicted single-pass transmembrane (TMEM) domain (Figure 1A). Domain analysis indicates that *MISTRAV* belongs to the poorly characterized B12D, NADH: ubiquinone reductase complex I MLRQ subunit family (pfam:06522). Using blastp analysis, we identified a 91 AA predicted ORF (SQPV78/YP_008658503.1) with high identity to human MISTRAV [47% (38/81) amino acid identity, 66% positives (54/81)](Figure 1B) in the squirrelpox genome, hereafter *vMISTRAV*.

**Figure 1.**
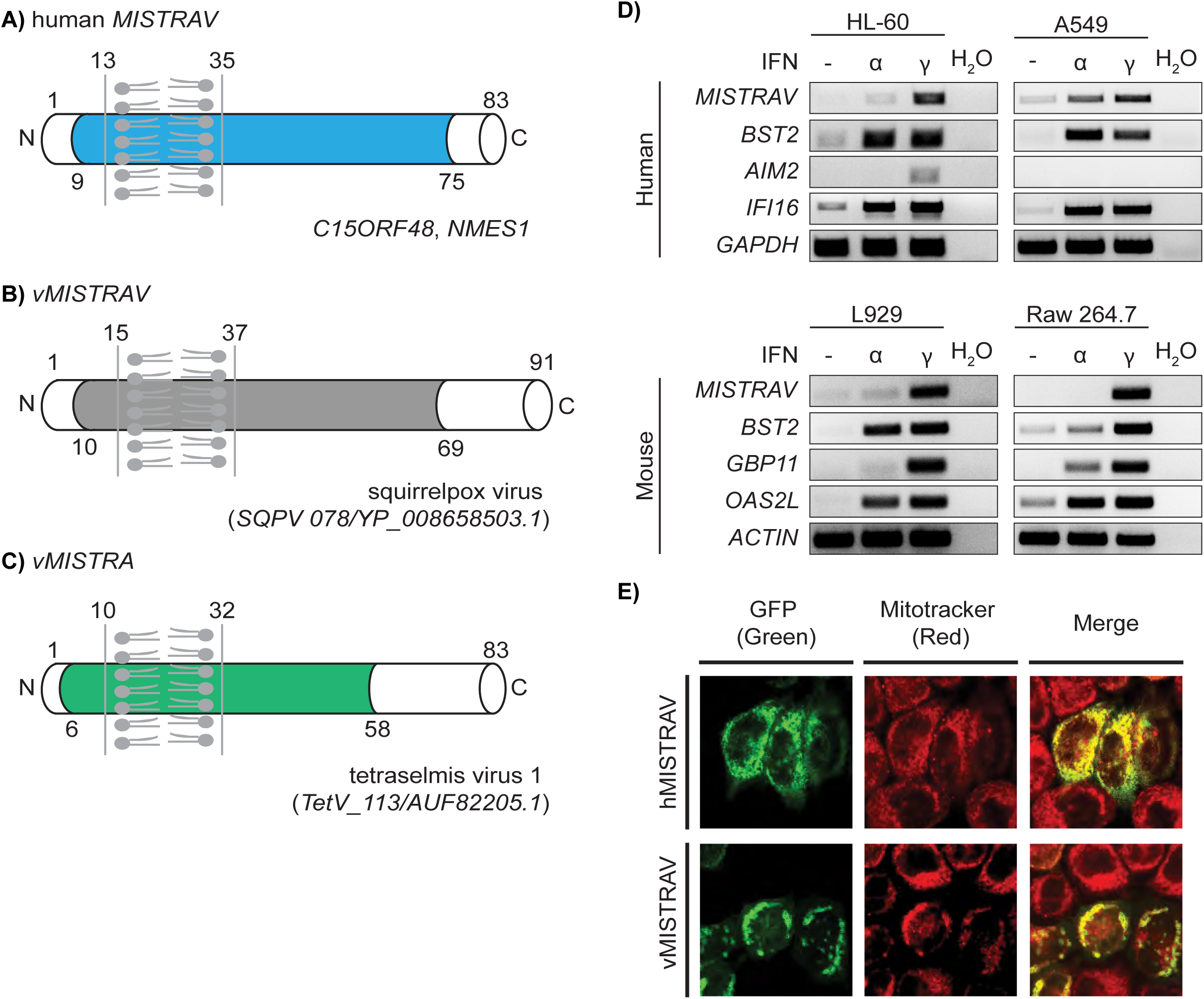
MISTRAV is a small interferon gamma-stimulated mitochondrial factor also encoded by divergent viruses. **A)** Diagram of *MISTRAV* with predicted domains indicated. Colored domain represents B12D, NADH: ubiquinone reductase complex I MLRQ subunit family (pfam:06522). Transmembrane predicted using TMHMM (http://www.cbs.dtu.dk/services/TMHMM/). **B)** Diagram of *vMISTRAV*, the *MISTRAV* homolog encoded by squirrelpox, annotated with predicted domains. **C)** Diagram of *vMISTRA*, the MISTR homolog identified in the genome of a giant virus which infects algae. **D)** RT-PCR using cDNA produced from RNA from interferon-treated human and mouse cell lines. *BST2*, *AIM2*, *IFI16*, *GBP11*, and *OAS2L* are interferon-stimulated gene (ISG) controls. **E)** Confocal images of HeLa cells transfected with constructs encoding *MISTRAV-* or *vMISTRAV-GFP*. *<u>MI</u>tochondrial <u>ST</u>ress <u>R</u>esponse <u>A</u>nti<u>V</u>iral* (*MISTRAV*), *<u>v</u>iral MISTRAV* (*vMISTRAV*), *<u>v</u>iral MISTR <u>A</u>lgae* (*vMISTRA*).

Reciprocal blastp analysis indicates *vMISTRAV* was presumably acquired by horizontal gene transfer derived from a host copy of *MISTRAV*. Specifically, using vMISTRAV AA sequence as a query returns numerous host MISTRAV sequences – and not sequences of MISTRAV paralogs - from diverse species (Additional details in Supplementary file 6). Consistently, domain analysis indicates vMISTRAV has a similar primary structure to host MISTRAV: short N-terminus, single-pass TMEM domain, longer C-terminus, and a B12D domain spanning these features (Figure 1B).

Subsequent database searches detected another *MISTR* ORF (*TetV-113/AUF82205.1*), hereafter *vMISTR <u>A</u>lgae* (*vMISTRA*), in the genome of the giant DNA virus (Schvarcz and Steward, 2018) Tetraselmis virus 1 (TetV-1) - a mimivirid - that infects the cosmopolitan green alga *Tetraselmis* (Figure 1C). *vMISTRA* encodes a predicted 83 AA ORF with a primary structure similar to MISTRAV and vMISTRAV: short N-terminus, predicted single-pass TMEM domain, longer C-terminal domain, and a B12D domain spanning these features. A clustal amino acid alignment using three *Tetraselmis* MISTR protein sequences from the database indicates that vMISTRA displays the greatest homology with A0A061RM32 in UniProt (40% identity by blastp) (Figure 1- figure supplement 1A-B). Thus, sequences resembling MISTR proteins are encoded by viruses that infect hosts from algae to mammals.

### *MISTRAV* is upregulated by interferon and localizes to the mitochondria

A hallmark shared by many immune defense factors critical to modulating infections is upregulation by immune signals such as interferon. To test whether *MISTRAV* is an ISG, we performed RT-PCR on samples from various human and mouse cell lines treated with either Type I (IFN-α) or Type II Interferon (IFN-γ). While *MISTRAV* is induced by IFN-α in A549 lung epithelial cells, our data indicate that *MISTRAV* is primarily upregulated by IFN-γ in the human and mouse cell lines tested (Figure 1D). Thus, *MISTRAV* displays two key hallmarks of crucial immune factors like OAS1: upregulation by immune signals and viral homologs (*vMISTRAV*, *vMISTRA*).

Both human and mouse (known as AA467197) MISTRAV have evidence for mitochondrial localization. The inventory of mammalian mitochondrial genes – MitoCarta (Calvo et al., 2016; Pagliarini et al., 2008) - detected MISTRAV in mitochondria across various tissues: small intestine, large intestine, stomach, placenta, and testis. In addition, MISTRAV is related to two known mitochondrial factors (MISTR1 and MISTRH) thought to be supernumerary factors associated with the electron transport chain (Balsa et al., 2012; Floyd et al., 2016; Tello et al., 2011). Consistent with previous findings, our expression of MISTRAV-GFP and vMISTRAV-GFP in HeLa cells revealed strong co-localization with the mitochondrial marker, MitoTracker (Figure 1E). Intriguingly, vMISTRAV-GFP expression in some cells resulted in altered morphology of the cell and/or mitochondria (Figure 1E).

### *MISTRAV* belongs to a gene family rapidly evolving in primates

*MISTRAV* and its poorly characterized paralogs *MISTR1* and *MISTRH* – are conserved over 450 million years of evolution as evidenced by the presence of orthologs in the zebrafish and spotted gar genomes (Figure 2- figure supplement 1). To gain insights into the recent evolution of all three MISTR proteins, we carried out evolutionary analysis using sequences for primate orthologs spanning more than 35 million years of divergence (Figure 2, Supplementary file 1, Supplementary file 6). Specifically, we tested if MISTR proteins display elevated rates of nonsynonymous amino acid substitution relative to synonymous substitution rates (dN/dS > 1) to determine if these proteins are likely to be engaged in genetic conflicts with pathogen-encoded inhibitors (McLaughlin and Malik, 2017) (Daugherty and Malik, 2012).

Comparative analyses of twenty-three primate orthologs using codon-based models implemented in PAML (Yang, 2007) (Figure 2, Supplementary file 6) revealed that both *MISTRAV* [M7 vs. M8 (F3X4) p < 0.0012] and *MISTR1* [M7 vs. M8 (F3X4) p < 0.0046] but not *MISTRH* [M7 vs. M8 (F3X4) p < 1.0000] display gene-wide rapid evolution patterns. Furthermore, these signatures in *MISTRAV* and *MISTR1* appear independent of any potential relaxed constraint within the predicted transmembrane (TMEM) domain as the signal is maintained when that domain is removed in additional tests [MISTRAV - M7 vs. M8 (F3X4) p < 0.0040, *MISTR1* - M7 vs. M8 (F3X4) p < 0.0040]. Calculating dN/dS values across the primate phylogeny using PAML identified multiple, distinct lineages in all three primate families [Hominoids (HOM), Old World Monkeys (OWM), and New World Monkeys (NWM)] with robust and recurrent signatures of rapid evolution for both *MISTRAV* and *MISTR1*.

Signatures of positive selection at specific amino acid residues can reveal key protein surfaces targeted by pathogen-encoded inhibitors and the number of surfaces with elevated dN/dS values is hypothesized to correlate with the number of interfaces (Daugherty and Malik, 2012). Using PAML (Yang, 2007), MEME (Murrell et al., 2012), FUBAR (Murrell et al., 2013)], we estimated dN/dS per amino acid site for *MISTR* genes. These analyses (Figure 2D, Figure 2E, Figure 2F) revealed seven different amino acid positions (∼8% of the whole-protein) distributed through *MISTRAV* with evidence of positive selection including two sites (21T and 79Q) identified by all three analyses. For *MISTR1*, three amino acid positions were predicted for rapid evolution owing to elevated dN/dS values, with 6I being notable for its detection by all three analyses.

**Figure 2:**
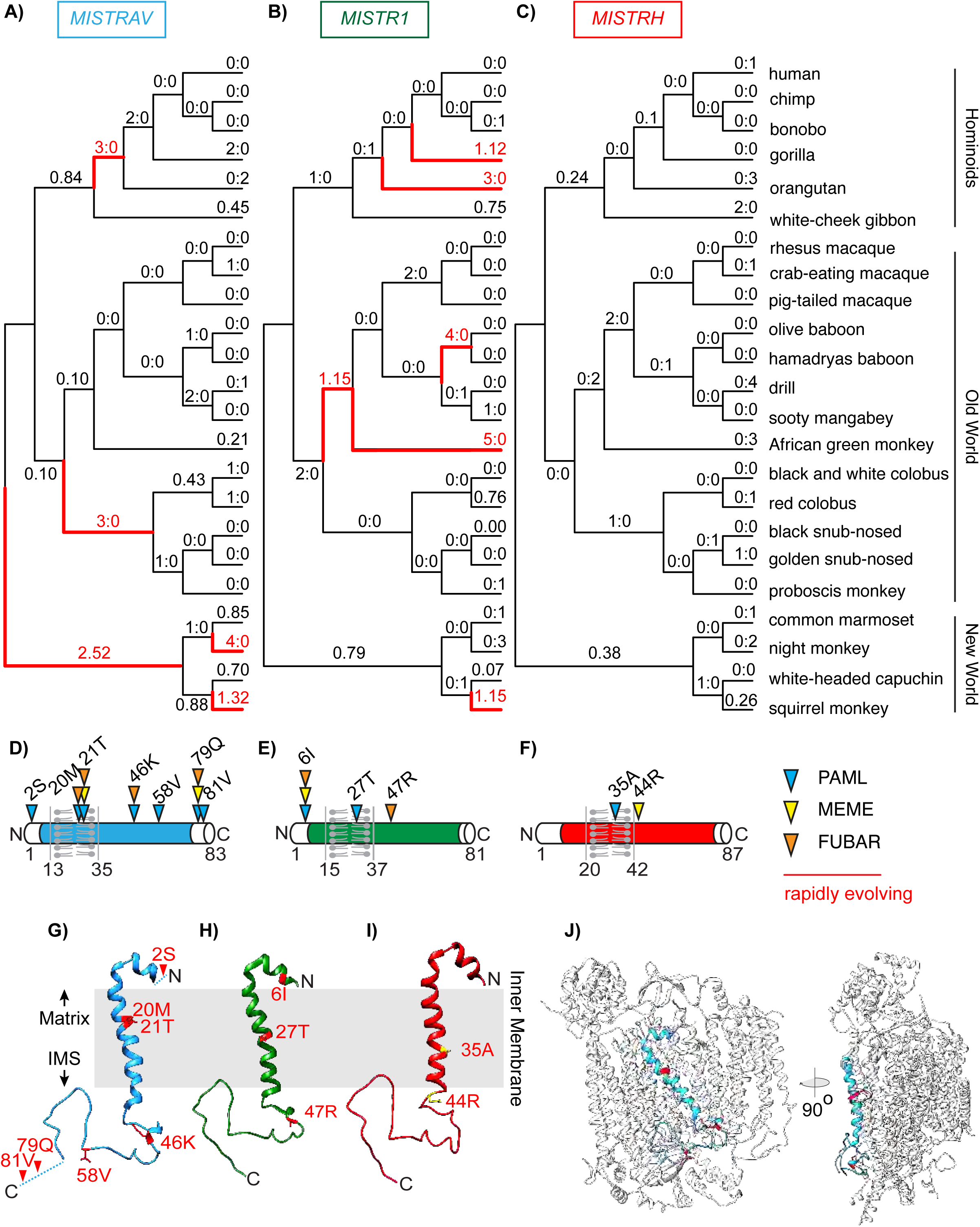
Rapid evolution of *MISTRAV* and its paralog *MISTR1* in primate genomes. Estimated dN/dS values predicted using FreeRatio analysis in PAML (Yang, 2007) across primate lineages for **A)** *MISTRAV*, **B)** *MISTR1*, and **C)** *MISTRH*. Rapidly evolving lineages (dN/dS > 1 or greater than or equal to 3 nonsynonymous amino acid substitutions: synonymous amino acid substitutions) are marked by red branches. **D)** *MISTRAV* **E)** *MISTR1*, and **F)** *MISTRH* amino acid positions predicted to be rapidly evolving (colored triangles) from PAML, MEME (Murrell et al., 2012), and FUBAR (Murrell et al., 2013) analysis. Numbering and residue are relative to the human reference sequence. Rapidly evolving sites for **G)** *MISTRAV* (red) **H)** *MISTR1* (red), and **I)** *MISTRH* (yellow) mapped onto the predicted structure of MISTR1. Models were generated using SWISS-MODEL (https://swissmodel.expasy.org/) based on the published structure of Complex IV of the electron transport chain containing MISTR1/NDUFA4 (PDB:5Z62)(Zong et al., 2018). **J)** Model of MISTRAV (blue) within Complex IV structure (silver).

Protein modeling with SWISS-MODEL (https://swissmodel.expasy.org/)(Waterhouse et al., 2018)(Figure 2G, Figure 2H, Figure 2I) using the only predicted structure of Complex IV to include MISTR1 [PDB:5Z62] (Zong et al., 2018) illustrates that MISTR TMEM domains are accessible for interfacing with cellular proteins. Thus, rapid evolution in the TMEM is unlikely to reflect relaxed constraint. Collectively, the rapid evolution signature observed for *MISTRAV* and *MISTR1* resemble that of other host factors that can dictate the outcomes of infections.

### Functional analyses support a role for *MISTRAV* and its encoded *miR-147b* in apoptosis

To investigate *MISTRAV* biology, we generated three A549 clonal cell lines – C15Δ1, C15Δ2, C15Δ3 - with distinct indels that disrupted the *MISTRAV* ORF using CRISPR/CAS (Figure 3A). A549 cells were selected because: 1) MISTRAV is interferon-inducible in these cells (Figure 1D, Figure 3B), 2) A549 cells are often used as a model for immune activation (Li et al., 2017), and relatedly, 3) this cell line is frequently used to model viral infections (Li et al., 2016) (e.g. coronaviruses, influenza, poxviruses). Consistent with the engineered mutations (Figure 3A), western blot analysis confirmed loss of MISTRAV protein in all three clones (Figure 3B). To maintain expression of a poorly characterized miRNA encoded by the 3’- UTR of *MISTRAV* (*miR-147b*) (Liu et al., 2009), we targeted the guide RNAs to exon 2 relative to the long *MISTRAV* isoform (Figure 3A, 875 nt) - a location where a frameshift in the RNA would be predicted to escape nonsense-mediated decay.

**Figure 3:**
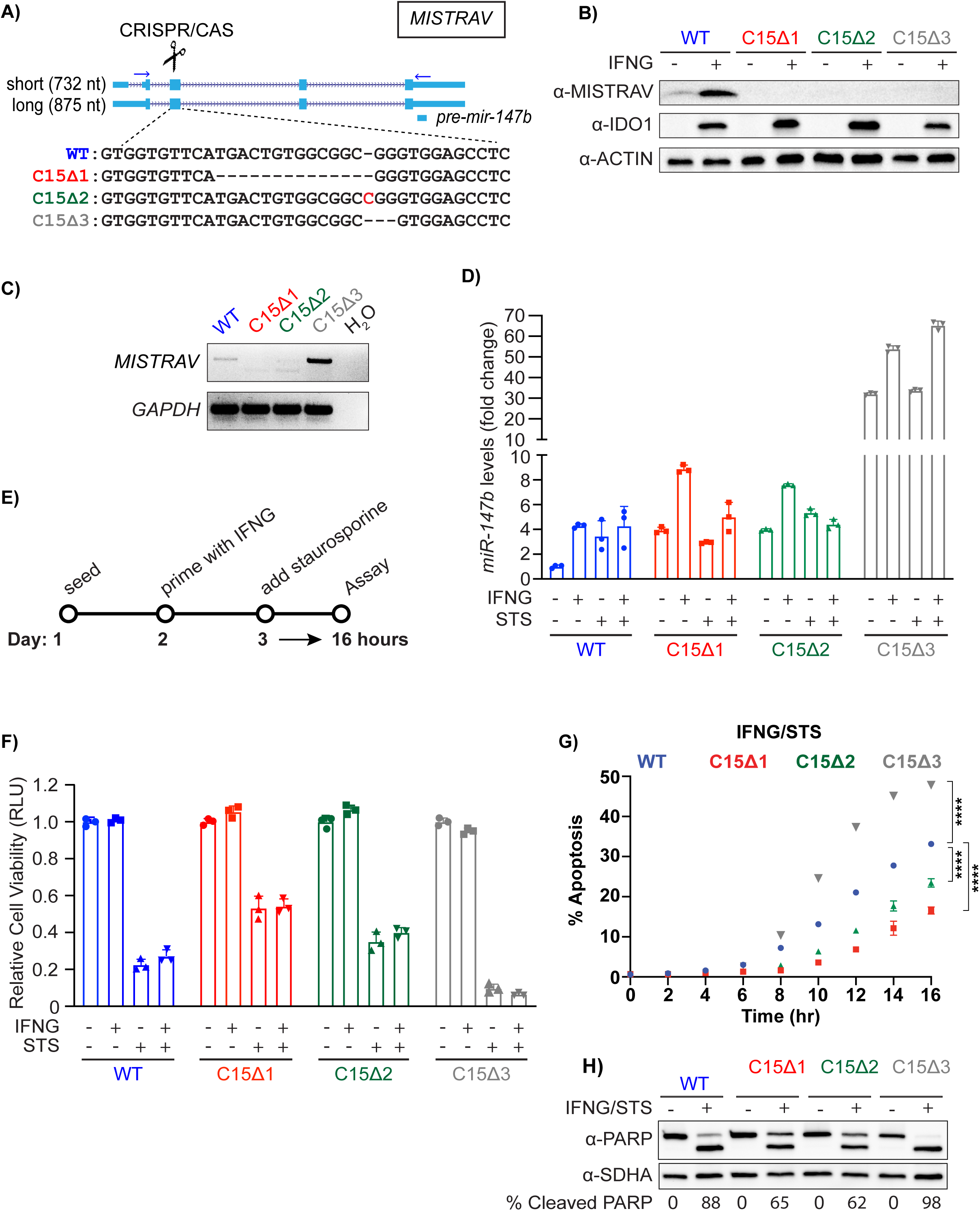
Loss-of-function analysis reveals a role for MISTRAV and its embedded miRNA – *miR- 147b* – in apoptosis. **A)** Diagram of the *MISTRAV* locus from the UCSC genome browser (http://genome.ucsc.edu/)(Kent et al., 2002). Two major transcripts are predicted for *MISTRAV*, which we term short (5 exons/predicted mRNA length 732 nt) and long (4 exons/predicted mRNA length 875 nt). The location of *pre-mir-147b* is marked by the blue box below predicted protein-coding mRNAs. Sequences of CRISPR-induced mutations targeted to exon 2 (relative to the long isoform of *MISTRAV*) in A549 cells, which result in predicted frameshifts. Deleted nucleotides are indicated by hypen (-) and inserted nucleotide is highlighted in red. **B)** Western blot analysis using protein lysates from IFN-γ treated A549 cells and *MISTRAV* deletion clones. IDO1 is an ISG control (Kane et al., 2016). **C)** RT-PCR analysis using primers (horizontal blue arrows) in **A)** on cDNA produced from total RNA extracted from IFN-γ- treated A549 WT and *MISTRAV* KO cells. **D)** *miR-147b* Taqman qPCR using RNA extracted from A549 WT and mutant cell lines treated with IFN-γ, STS, or both for 16 hours. *miR-423* was used as the endogenous control. Fold changes in *miR-147b* levels are relative to the *miR-147b* level in WT untreated cells. Data represent means ± SD (n= 3 replicates). **E)** Experimental timeline of apoptosis assays using WT and mutant A549 cells. **F)** CellTiter Glo (luciferase-based) cell viability assay on WT and mutant cells treated with IFN-γ, STS, or both for 16 hours. Data represent means ± SD (n= 3 replicates). **G)** Percent apoptosis of A549 WT and *MISTRAV* KO cells pre-treated with IFN-γ for twenty-four hours followed by STS treatment for 16 hours; Caspase 3/7 activity was normalized to the number of cells at the initial treatment timepoint measured by IncuCyte. Data represent means ± SD (n= 3 replicates). Statistical significance was determined by a two-tailed unpaired t-test, ****p*≤*0.0001. **H)** Western blot analysis of cleaved PARP in WT and *MISTRAV* KO cells treated with IFN-γ and STS. SDHA serves as loading control. Densitometry analysis of PARP levels was performed using Image Lab version 6.0.1 (Bio-Rad). % Cleaved PARP= (cleaved PARP/(Full + Cleaved PARP)) * 100. IFN-γ – interferon gamma, STS – staurosporine.

RT-PCR indicated that C15Δ1 and C15Δ2 cells lack full-length (FL) *MISTRAV* RNA expression in IFN-γ treated cells at steady-state while C15Δ3 cells display a fortuitous and drastic increase of the same transcript (Figure 3C). miRNA qPCR demonstrated that C15Δ1 and C15Δ2, maintain *miR-147b* at levels comparable to wild-type with expression of *miR-147b* in C15Δ3 ∼30 fold greater than WT (Figure 3D). Thus, C15Δ1 and C15Δ2 lack MISTRAV protein but maintain the miRNA, while C15Δ3 lacks MISTRAV with a gain-of-function of *miR-147b* expression.

Based on *MISTRAV* mitochondrial localization and numerous documented connections between immune responses involving cell death mediated through mitochondria, we reasoned that MISTR might mediate apoptotic responses. We primed WT and KO cells with IFN-γ then added the commonly used activator of apoptosis, staurosporine (STS), for 16 hours followed by functional analysis (Figure 3E). Assays were normalized to either untreated controls or to the number of cells being tested to account for differences in proliferation rates (Figure 3- figure supplement 1). Interestingly, we observed that C15Δ1 and C15Δ2 displayed reduced sensitivity to STS, while C15Δ3 showed increased sensitivity to STS compared to WT cells (Figure 3F). Consistent with these results, we detected robust caspase-3/7 cleavage activity for C15Δ3 compared to WT (Figure 3G; p<0.0001). In addition, detectable decreases in caspase-3/7 activity were observed for C15Δ1 and C15Δ2 relative to WT treated cells (Figure 3G; p<0.0001). The differential sensitivities of the clones to STS relative to WT were consistent with levels of PARP cleavage across the clones in response to STS (Figure 3H). These data suggest a role for MISTRAV and *miR-147b* in apoptosis.

### Ultraconserved miRNAs link MISTR paralogs

To gain insights into the increased levels of apoptosis in C15Δ3 cells associated with *miR-*147b [*miR-147* in mouse (Liu et al., 2009)] we performed comparative miRNA target analysis. A recent survey indicates that the *miR-147b* seed sequence is conserved in vertebrate orthologs (Bartel, 2018). Strikingly, our sequence analysis demonstrated that all twenty-two nucleotides of *miR-147b* miRNA are identical between human and spotted gar; which represents around 450 million years of divergence from a common ancestor (Figure 4). Interestingly, although the *MISTRAV* locus is present in the zebrafish genome, *miR-147b* sequence is likely non-functional because of disruptive indels (Figure 4, Figure 4- figure supplement 1).

**Figure 4:**
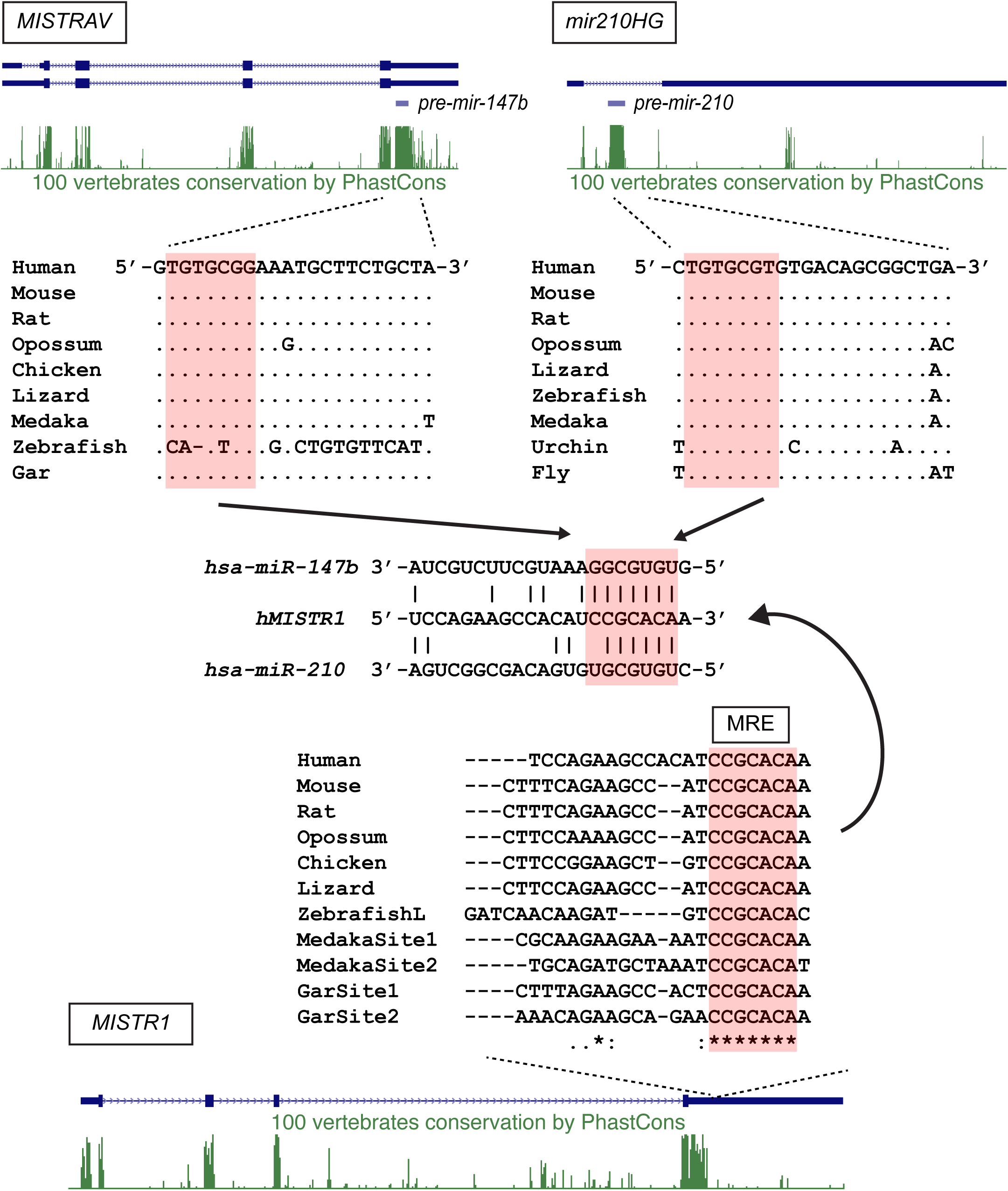
Ultraconserved miRNAs are predicted to target a vertebrate-specific miRNA response element in *MISTR1*. Human *MISTRAV*, *mir210HG*, and *MISTR1* loci with predicted gene structures and PhastCons [green peaks (Siepel et al., 2005)] track from the UCSC genome browser are shown. Orthologous sequences were retrieved from the NCBI sequence database (Supplementary file 6). Predicted seeds and miRNA response element (MRE) are marked by salmon-colored boxes.

miRNA target analysis uncovered 36 [(mirdb.org)(Wong and Wang, 2015)(Liu and Wang, 2019)] and 19 [Targetscan (www.targetscan.org) (Agarwal et al., 2015)] *miR-147b* predicted targets (Supplementary files 2 and 3), of which only two were shared by both databases: *C11orf87* and the *MISTRAV* paralog, *MISTR1*. The predicted miRNA response elements (MRE) in the 3’-UTR of the *MISTRAV* paralog, *MISTR1*, is a predicted 8mer seed that is perfectly conserved out to fish genomes (Figure 4). In addition, 1) the 8mer has duplicated in some fish *MISTR1* orthologs (e.g. gar and medaka)(Figure 4) and 2) zebrafish maintains the predicted MRE for *miR-147b*.

Interestingly, the predicted MRE encoded by *MISTR1* overlaps with an MRE for an unrelated miRNA, *miR-210*. *miR-210* is highly upregulated by HIF1α during low oxygen conditions and thought to be critical for the hypoxic response (Huang et al., 2009). Assays using a MRE reporter encoding the human *MISTR1* (*NDUFA4*) 3’-UTR (Bertero et al., 2012) support the functionality of this shared MRE, yet the significance has remained an open question. Evolutionary analysis indicates that the *miR-210* seed is perfectly conserved in bilateria for sequences sampled, with 19/22 nucleotides identical between the human and *Drosophila* orthologs (Figure 4) and 21/22 nts identical between human and fish orthologs. Thus, the *MISTR1* 3’-UTR encodes a highly conserved MRE potentially targeted by two distinct ultraconserved miRNAs with an overlapping seed sequence; one of which is encoded by the paralog *MISTRAV*.

### MISTR1 is regulated by stress-inducible miRNAs

TargetScan predicts seven MREs in the *MISTR1* 3’-UTR for six distinct miRNAs [*miR-7-5p*, *miR- 145-5p* (2 sites), *miR-147b-3p*, *miR-202-5p*, *miR-205-5p* and *miR-210-3p*], which have seed sequences that are highly conserved in vertebrates with a subset extending in sequence conservation to bilateria (Figure 5A)(Bartel, 2018). MRE reporter assays using a luciferase reporter with the entire 1685 bp human *MISTR1* 3’-UTR (Figure 5B) revealed that transient co-transfection of either *miR-7-5p*, *miR-147b-3p*, *miR-210-3p* in HEK293T cells resulted in dramatic knockdown (40-65% of vector alone). Correspondingly, western blots with lysates from HEK293T and A549 cells transiently transfected with *miR-7-5p*, *miR-147b-3p*, *miR-210-3p* (Figure 5C) demonstrated knockdown of endogenous MISTR1 protein. We identified two polyA signal canonical hexamers (AATAAA; 161-166, 1666-1671 relative to human 3’-UTR) in the *MISTR1* 3’-UTR that divides the first four MREs from the three downstream sites (Figure 5A). Interestingly, the miRNAs that did not result in knockdown are located downstream of the first polyA signal while those that did cause knockdown of targets are located upstream of the first polyA signal. Therefore, the *MISTR1* 3’-UTR encodes several predicted MREs for conserved miRNAs, of which a subset is functional in cell culture assays.

**Figure 5.**
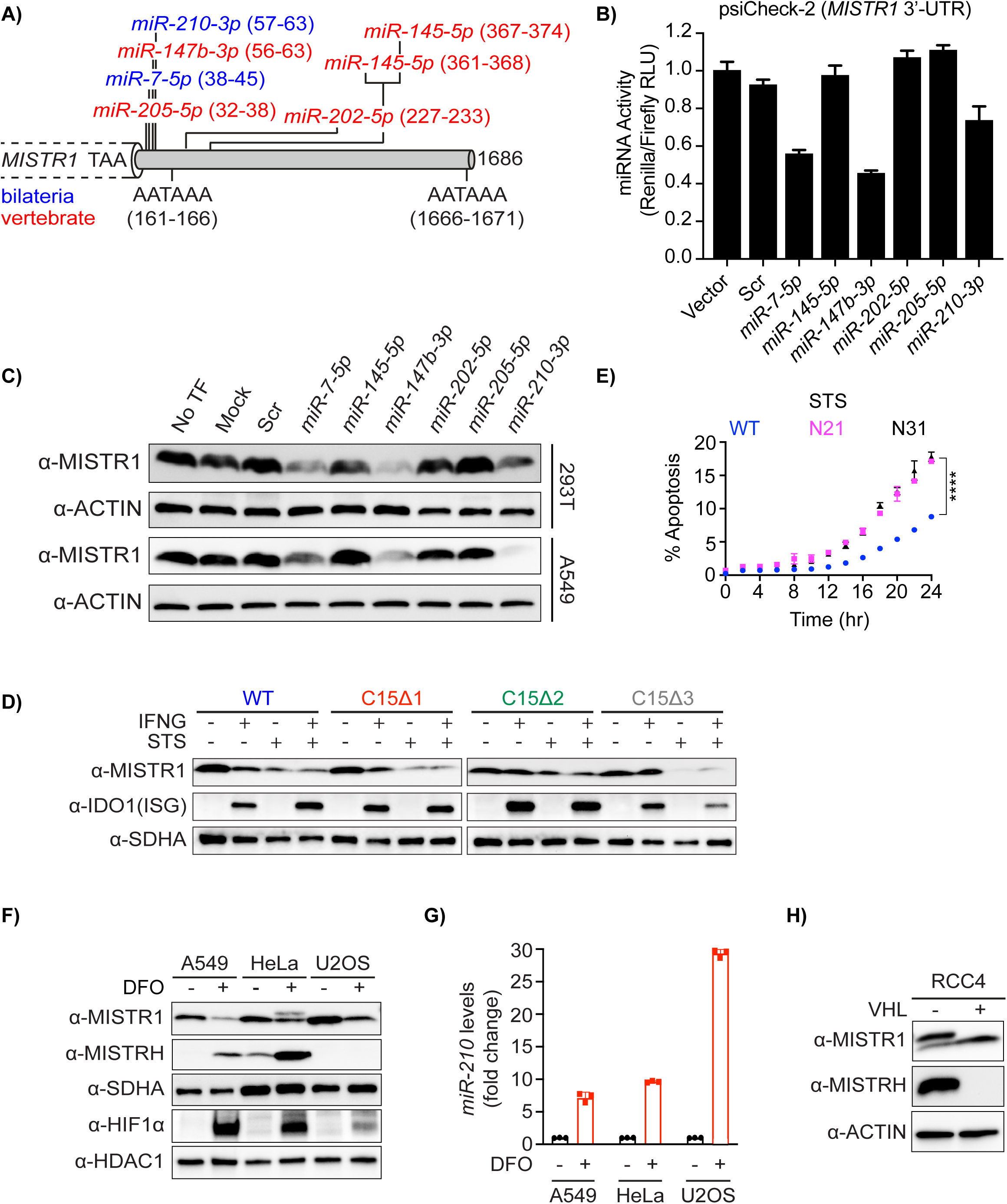
MISTR1 is a target of multiple conserved miRNAs, ubiquitously expressed, and downregulated by stress. **A)** Diagram of predicted MREs in the full-length human *MISTR1* 3’-UTR. Numbering is relative to the first nucleotide downstream of the stop codon for the *MISTR1* human reference sequence. MREs are colored by miRNA seed conservation determined by (Bartel, 2018): bilateria (blue) and vertebrate (red). Identified core polyA signal sequence motifs (5’-AATAAA-3’) are highlighted. **B)** miRNA reporter assays for miRNAs predicted to target *MISTR1*. psiCheck2 encoding the full-length human *MISTR1* 3’-UTR and candidate miRNAs were sequentially transfected into HEK293T cells followed by luciferase assays. Data represent means ± SD (n= 3 replicates). **C)** Western blot for endogenous MISTR1 levels in HEK293T and A549 using lysates from cells transfected with miRNAs predicted to bind the *MISTR1* 3’-UTR. **D)** Western blot for endogenous MISTR1 levels using lysates from A549 WT or *MISTRAV* KO cells treated with IFN-γ, STS, or both for 16 hours. α-SDHA blot serves as a control for mitochondrial protein stability. **E)** Percent apoptosis of A549 WT and *MISTR1* KO cells treated with STS; Caspase 3/7 activity was normalized to the number of cells at the initial treatment timepoint measured by IncuCyte. Data represent means ± SD (n= 3 replicates). Statistical significance was determined by a two-tailed unpaired t-test, ****p*≤*0.0001. **F)** Western blot analysis of MISTR1 and MISTRH levels 24 hours after chemical hypoxia induction by DFO. The upper band in the DFO-treated HeLa lane in the α-MISTR1 blot is MISTRH. SDHA – mitochondrial control, nuclear HIF1*α −* hypoxia control, HDAC1 – nuclear protein control. **G)** *miR-210* Taqman qPCR of cell lines in **F)** following 24 hours of DFO treatment. *miR-423* was used as the endogenous control. Fold changes in *miR-210* levels in DFO-treated cells are relative to the *miR-210* level in untreated cells. Data represent means ± SD (n= 3 replicates). **H)** Western blot analysis of MISTR1 and MISTRH levels in the RCC4 kidney cancer cell line with or without stable rescue expression of Von Hippel Lindau (VHL). The upper band in the RCC4 (- VHL) lane in the α-MISTR1 blot is MISTRH. IFN-γ – interferon gamma, STS – staurosporine, DFO – deferoxamine mesylate.

To test if MISTR1 is downregulated by stress, we performed western blots on lysates from A549 WT and *MISTRAV* A549 KO cells treated with STS and/or interferon. We observed a progressive downregulation of MISTR1 following treatment with STS or STS/IFN-γ compared to IFN-γ alone (Figure 5D). We also observed a nearly complete loss of MISTR1 in C15Δ3 mutant cells, which overexpress *miR-147b*. MISTR1 downregulation appears either specific or rapid in comparison to levels of the mitochondrial Complex II protein SDHA, which are largely unchanged under the same conditions (Figure 5D). These data suggest loss of MISTR1 may promote apoptosis under conditions of stress. To test this hypothesis directly, we generated two MISTR1 KO A549 clonal cell lines (N21 and N31)(Figure 5, figure supplement 1). A control experiment showed that MISTR1 KO cells exhibit rates of proliferation similar to WT (Figure 5- figure supplement 1D). Assay of these cells following STS treatment using live-cell analysis with the IncuCyte and western blot for cleaved PARP indicates that *MISTR1* KO cells are more sensitive to this apoptotic trigger compared to WT cells (Figure 5E, Figure 5- figure supplement 1E).

Next, we examined regulation of MISTR1 in cells under hypoxic stress; a condition when *miR-210* and MISTR1’s paralog, MISTRH (Tello et al., 2011), are expressed. Following the induction of chemical hypoxia by deferoxamine mesylate (DFO) treatment in three cell lines, we observed downregulation of MISTR1 concomitant with an upregulation of HIF1α, MISTRH (Figure 5F), and *miR-210* (Figure 5G). Analysis of RCC4 kidney cancer cells with and without Von Hippel-Lindau (VHL) tumor suppressor indicate that the opposing expression of MISTR1 and MISTRH requires HIF signaling (Figure 5H). Thus, MISTR1 is downregulated by ultraconserved stress-induced miRNAs under conditions when its paralogs are upregulated.

### A broad phylogenetic distribution of MISTR proteins

To examine the implications of our findings in an evolutionary context, we characterized the breadth of MISTR proteins across eukaryotic genomes. While a recent study detected *MISTR1* homologs in yeasts, including Baker’s and fission yeast, as well *Plasmodium* (Balsa et al., 2012), major gaps in the distribution and evolution of these proteins remain. We identified additional predicted proteins across animals and plants displaying homology to MISTR variants (Supplementary file 4). These data indicate *MISTRAV*, *MISTR1*, *MISTRH* sequences are conserved in vertebrate genomes with duplications present in the zebrafish genome for *MISTR1* and *MISTRH*; a phenomenon common to genes of teleost fish (Howe et al., 2013).

Maximum-likelihood phylogenetic analysis using PhyML (Guindon et al., 2010) of these AA sequences defines three major clades: A) vertebrate MISTRAVs, B) vertebrate MISTR1 and MISTRHs as well as Nematostella and *Drosophila* proteins, and C) plant MISTRs along with algae and yeast proteins (Balsa et al., 2012). Low bootstrap values observed throughout the tree may be a consequence of the small length of MISTR sequences (*i.e.* too few characters) despite generating one-thousand trees for bootstrap analysis.

The clustering of vMISTRAV between the clade representing MISTRAV from mammals and lineages leading to chicken and zebrafish support the notion that *vMISTRAV* 1) likely originated from a mammalian host in agreement with the primary host of squirrelpox (Darby et al., 2014) and diverged substantially after horizontal gene transfer and 2) is derived from host *MISTRAV* and not *MISTR1* or *MISTRH*. A similar placement of *vMISTRA* from TetV-1 near MISTR sequence from the *Tetraselmis* algae protein (A0A061RM32) is also consistent with horizontal transfer from host to virus. Interestingly, the choanoflagellate *Salpingoeca rosetta* encodes two divergent MISTR homologs as evidenced by XP_004989268.1 clustering with Clade B and XP_004998377.1 clustering with Clade C. These data indicate that MISTR is widely distributed in genomes of diverse eukaryotes and has undergone repeated diversification, including ancestral duplications, as well as more recent evolutionary innovations.

### vMISTRAV antagonizes apoptotic responses

Our data indicate a role for MISTR in cellular stress responses. To test the ability of vMISTRAV to counteract these responses, we engineered cells stably expressing the squirrelpox protein with a C-terminal HA epitope tag (Figure 7A). vMISTRAV-expressing cells grow at the same rate as control cells expressing an empty vector (EV) (Figure 7- figure supplement 1A). When vMISTRAV cells were treated with three activators of apoptosis – STS, actinomycin D (ActD), and camptothecin (CPT), we observed a protective effect of vMISTRAV as indicated by marked decreases in Caspase 3/7 activity (% apoptosis) (Figure 7C, Figure 7D, Figure 7E) as well as decreases in percentage of cleaved PARP (Figure 7- figure supplement 1B) compared to EV controls. We therefore conclude that the virus-encoded vMISTRAV inhibits apoptosis triggered by distinct mechanisms, consistent with a newly described host-pathogen conflict for control over the persistence of virus-infected cells.

**Figure 6:**
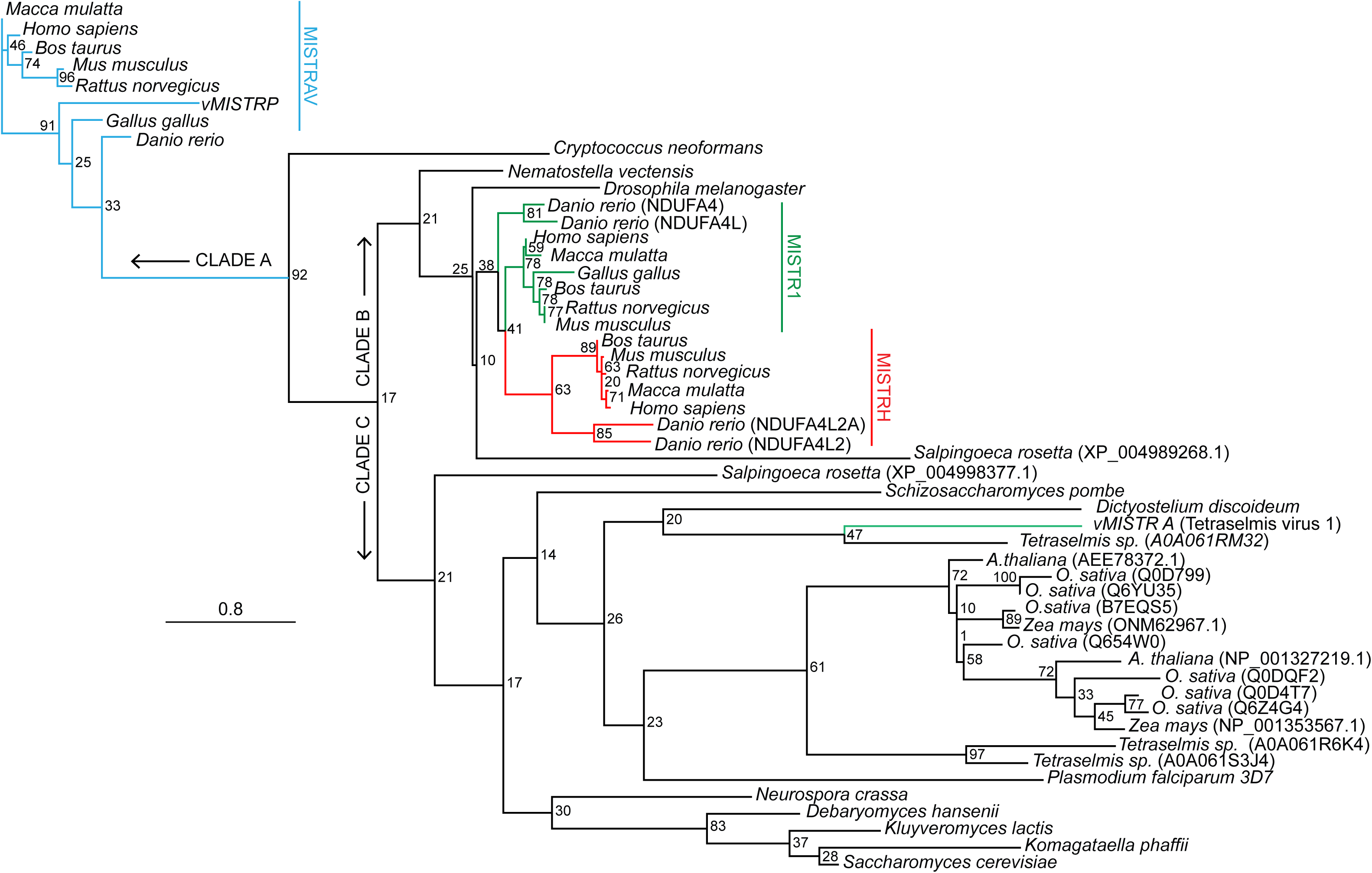
A broad phylogenetic distribution of MISTR sequences. An inferred tree built using MISTR amino acid sequences by maximum-likelihood analysis using PhyML (Guindon et al., 2010) (http://www.atgc-montpellier.fr/phyml/) with the LG +G model and 1000 bootstraps. Sequences were extracted from the NCBI sequence database, Uniprot (https://www.uniprot.org/) and (Balsa et al., 2012) (Supplementary file 4). Bootstrap percentages from analysis are placed at nodes. Scale for amino acid substitutions per site – bottom left.

**Figure 7:**
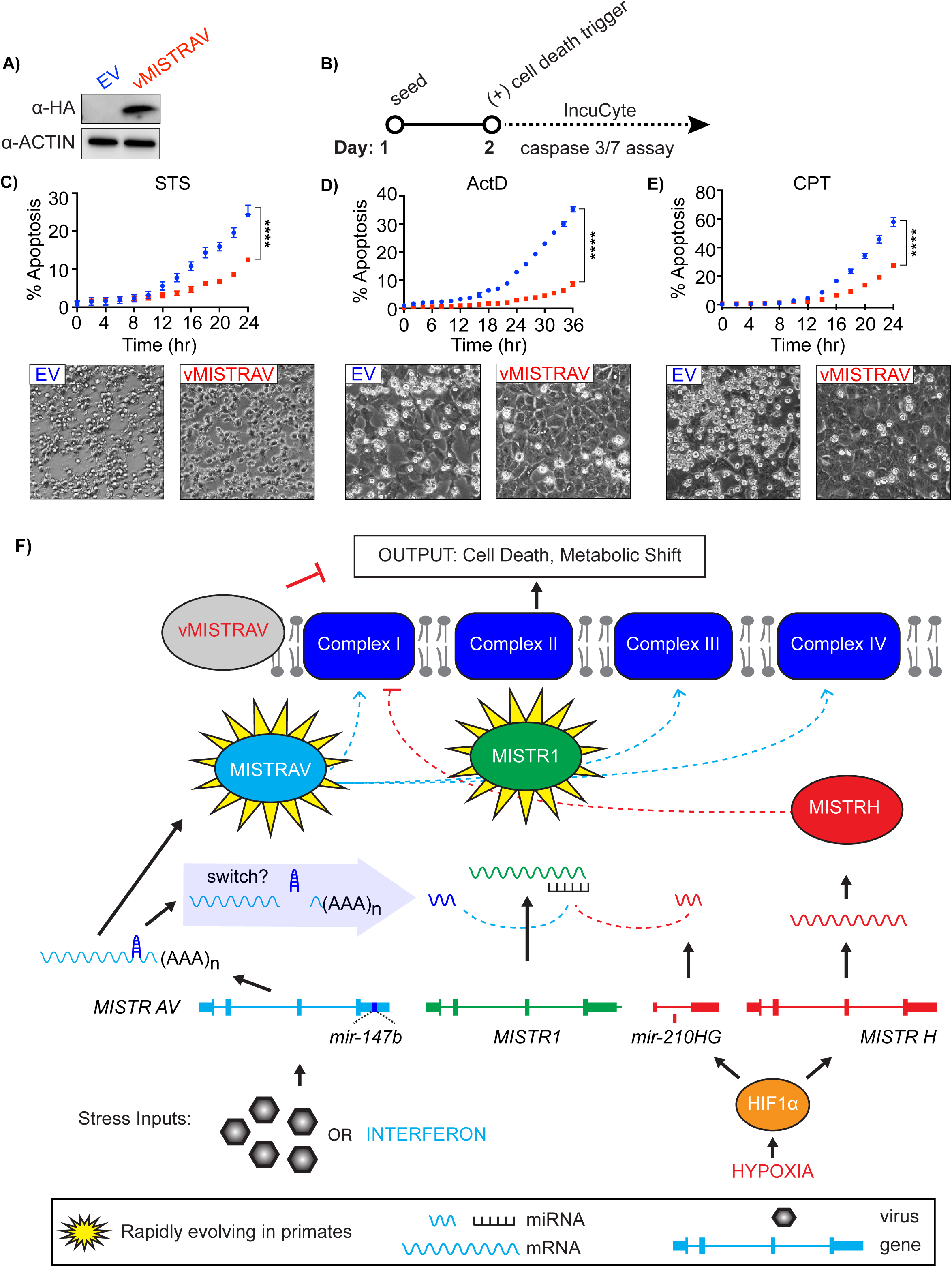
<u>MI</u>tochondrial <u>ST</u>ress <u>R</u>esponse Circuit (MISTR) Model. **A)** Western blot for vMISTRAV-HA using lysates from stably-expressing cells. EV – empty vector. **B)** Apoptosis assay timeline for vMISTRAV cells. Percent apoptosis and phase contrast images of EV and vMISTRAV-expressing cells following treatment with distinct activators of apoptosis: **C)** staurosporine (STS), **D)** actinomycin D (ActD), and **E)** camptothecin (CPT). Phase contrast images were taken 16 hours after treatment with STS or ActD and 24 hours after treatment with CPT. Caspase 3/7 activity was normalized to the number of cells at the initial treatment timepoint measured by IncuCyte. Percent apoptosis data represent means ± SD (n= 4 replicates). Statistical significance was determined by a two-tailed unpaired t-test, ****p*≤*0.0001 **F)** Schematic diagram of a MISTR network shows proposed interactions between the electron transport chain complexes and vertebrate MISTR proteins (MISTRAV, MISTR1, MISTRH, vMISTRAV). MISTR loci and RNA produced from them including *miR-147b* and *miR-210* are also illustrated. Using the example of infection as a stressor, *MISTRAV* transcription is induced by interferon (Figure 1D, Figure 3B) resulting in the production of MISTRAV RNA. MISTRAV protein localizes to the mitochondria (Figure 1E), to promote host defense. In the model, *miR-147b* production acts to inactivate MISTRAV translation and downregulate MISTR1 to facilitate the apoptotic response (Figure 3 and Figure 5). Virus encoded variants (vMISTRAV) counteract the response by inhibiting apoptosis through resemblance to MISTR components. In the case of hypoxic stress, *MISTRH* and *mir-210HG* are transcribed from distinct loci to produce MISTRH (Tello et al., 2011), which can inhibit Complex I activity (dashed red line), while *miR-210* (Huang et al., 2009) downregulates MISTR1 to facilitate the cellular hypoxic response. Rapid evolution of *MISTRAV* and *MISTR1* (Figure 2) is highlighted by yellow stars. Blue dashed lines from MISTRAV indicate potential ETC complex interactions from published data including protein-protein interactions proposed from mass spec analysis (Floyd et al., 2016). Although MISTR proteins may be embedded components in the mitochondrial inner membrane undergoing stress regulated and miRNA-mediated exchanges, they are shown as circles for clarity in the model.

## DISCUSSION

### *MISTRAV* displays hallmarks of a critical immune defense function

Here we characterized a combination of features common to crucial immune factors to discover how conserved, but mostly uncharacterized cellular proteins can mediate the key host defense process of apoptosis. It is known that a subset of ISGs provide critical defenses against invading pathogens (Schneider et al., 2014; Schoggins, 2014). However, of the more than 400 ISGs identified to date, the majority are poorly characterized (Schoggins et al., 2011; 2014). Therefore, a high priority of immunological research is to assign functions, define interactions, and uncover regulatory mechanisms for this collection of vital gene products.

We define *MISTRAV* as an IFN-γ-inducible gene (Figure 1D) and protein (Figure 3B), which builds on previous work showing that *MISTRAV* is induced by other immune signals: LPS, poly I:C, and PAM3SCK4 in primary mouse and human macrophage cell lines (Liu et al., 2009), LPS in human primary effector dendritic cells (Zimmer et al., 2012), and IFN-α (Schoggins et al., 2011). Several lines of evidence suggest that cellular MISTRAV is targeted for inactivation by multiple pathogens. Specifically, signatures of rapid evolution we detected in primate genomes for *MISTRAV* (Figure 2A, Figure 2D) point to repeated antagonistic interactions with pathogens on multiple protein surfaces. Although the precise functions of the three MISTRAV domains remain undefined (N-terminus, transmembrane domain, and C-terminus), all display evolutionary patterns consistent with genetic conflicts (Barber and Elde, 2014)(Elde et al., 2008; Sawyer et al., 2004). We predict that rapidly evolving surfaces on opposite sides of the TMEM, which may be otherwise shielded by the mitochondrial inner membrane, represent unique surfaces targeted by pathogen-encoded inhibitors.

While positive selection predicts direct inhibitors of MISTRAV and MISTR1 functions, the presence of two viral homologs (*vMISTRAV* and *vMISTRA*) supports the idea that viruses also counteract this defense pathway via mimicry. Independent acquisition of related proteins by viruses that infect highly divergent hosts appears to be extremely rare with the largest evolutionary span thus far being distinct copies of IL-10 encoded by herpesviruses which infect fish and mammals (Ouyang et al., 2014). To our knowledge, these are the first ETC-associated genes known to be acquired by viruses. These observations indicate that the MISTR pathway provides a vital cellular defense that can influence the outcome of infections. Consistent with this idea, we demonstrate the ability of vMISTRAV to curb apoptotic responses from stimuli that function by distinct mechanisms (Figure 7). Notably, while *MISTRAV*, *OAS1* (Hancks et al., 2015; Mozzi et al., 2015), *cGAS*, *MX1* (Mitchell et al., 2012), *APOBEC3G* (Sawyer et al., 2004), *ZAP* (Daugherty et al., 2014; Kerns et al., 2008), *BST* (tetherin) (Lim et al., 2010; McNatt et al., 2009) and *PKR* (Elde et al., 2008) are all rapidly evolving and upregulated by interferon, only MISTRAV and OAS1 (Darby et al., 2014) homologs are known to be encoded in virus genomes.

### MISTR1 bridges the electron transport chain and stress responses

MISTR1 has been shown to associate with ETC complexes and was presumed to act as a structural component of the complex, but additional functional roles are a matter of debate (Balsa et al., 2012; Kadenbach, 2017; Pitceathly and Taanman, 2018; Pitceathly et al., 2013). MISTR1 loss-of-function caused by a homozygous splice donor mutation is associated with the neurological disorder Leigh’s syndrome (Pitceathly et al., 2013). MISTR1’s annotation as NDUFA4 comes from initial findings that it co-purifies with Complex I (Carroll et al., 2006). More recent work provided evidence for a primary Complex IV association (Balsa et al., 2012). The presence of MISTR1 on the external surface (Figure 2J) of Complex IV was interpreted as a means of regulating higher-order ETC complex formation into supercomplexes (Zong et al., 2018). Our data implicate downregulation of MISTR1 as a critical step for cells to respond to stresses, including pathogen infections (Figure 5). High levels of conservation of *MISTR1* MREs for *miR-210* and *miR-147b* (Figure 4) suggest the necessity of downregulating MISTR1 during immune signaling and hypoxia (Figure 5).

### MISTR is a vertebrate specific stress response circuit

Integrating evolutionary analysis with experimental genetics and related functional analysis led us to define a model for the <u>MI</u>tochondrial <u>ST</u>ress <u>R</u>esponse circuit (MISTR)(Figure 7F). While some previous studies hinted at potential interactions for MISTR components, functional connections were largely unknown. For instance, *miR-147b* and *miR-210* were shown to share a seed sequence and these miRNAs can downregulate a MRE reporter encoding the *MISTR1* 3’-UTR when transfected (Bertero et al., 2012). In addition, *miR-147b* functions were recently associated with the TCA cycle (Zhang et al., 2019), but the observation that *miR-147b* was encoded by the same gene as a *MISTR1* paralog had not been reported (Bertero et al., 2012)(Figure 4). Likewise, the overexpression of endogenous MISTRH correlating with loss of MISTR1 protein (Figure 5H) has been observed in <u>c</u>lear <u>c</u>ell <u>R</u>enal <u>C</u>ell <u>C</u>arcinoma (ccRCC) tumor samples and ccRCC cell lines (Minton et al., 2016); a disease characterized by hyperactive HIF signaling (Brugarolas, 2014), but the requirement of HIF in the regulation of this newly proposed circuit had not been tested (Figure 5H).

Our model predicts that MISTR1 is a ubiquitously expressed sensor of stress. Specific stress signals induce miRNA expression leading to the downregulation of MISTR1 and its replacement by inducible paralogs to facilitate apoptosis or some form of stress tolerance. Striking conservation of the miRNAs targeting *MISTR1* and cognate MREs (Figure 4) indicate MISTR-like responses are likely common in many diverse vertebrate species. In sharp contrast, MISTRAV is rapidly evolving near the C-terminus approximately only 80-100 bases upstream of the ultraconserved *miR-147b*.

The embedded nature of *miR-147b* implies a step-wise molecular progression of this response. Specifically, processing of *miR-147b* from the *MISTRAV* RNA, in principle, could uncouple the mRNA cap from the polyA tail rendering translation of MISTRAV infeasible. Consistent with this prediction and our findings, MISTRAV and *miR-147b* likely have related but separate functions (Figure 3). Furthermore, post-transcriptional mechanisms might also regulate mature *miR-147b* activity or its ability to target *MISTR1*. Strikingly, despite the high levels of *miR-147b* in C15Δ3 (Figure 3D), including at baseline, gross downregulation of MISTR1 - associated with the gain-of-function mutation in C15Δ3 - does not occur until STS is present (Figure 5D).

In contrast to *MISTRAV/miR-147b*, *miR-210* and *MISTRH* are encoded at distinct loci in an arrangement more permissive to complementary functions. However, *miR-210* is located within an intron of an uncharacterized non-coding RNA – called *miR-210HG* in humans. Here, processing of *miR-210* would not be predicted to inactivate the host gene. Therefore, *miR-210* and *miR-210HG* may share currently uncharacterized complementary functions. The distinct arrangements of *miR-147b* and *miR-210* are consistent with differences in cellular responses to hypoxia and infection. Namely, under hypoxia the cell will buffer itself from low-oxygen conditions enabling survival, while during infections there are more drastic, escalating levels of responses culminating in apoptosis to eliminate virus infected cells. Putting these findings together, the <u>MI</u>tochondrial <u>ST</u>ress <u>R</u>esponse (MISTR) system represents an evolutionarily dynamic circuit interfacing with fundamental cellular processes to mediate stress responses that can be targeted by viruses.

## MATERIALS AND METHODS

### Sequence analysis

Domain searches were performed using Interpro (https://www.ebi.ac.uk/interpro/), NCBI Conserved Domains (https://www.ncbi.nlm.nih.gov/Structure/cdd/wrpsb.cgi), and TMHMM for transmembrane domain prediction (http://www.cbs.dtu.dk/services/TMHMM/).

### Rapid evolution analysis

Primate nucleotide sequences were retrieved from the NCBI database (Supplementary file 1, Supplementary file 6). Multiple sequence alignments (MSA) were performed using Muscle in Geneious 11.1.5 (BioMatters). Indels were removed from alignments by manual trimming. To obtain dN/dS lineage estimates, the MSA for each gene and newick phylogenetic tree of sampled primates [based on known relationships (Perelman et al., 2011)] served as input for FreeRatio analysis implemented in PAML (Yang, 2007). PAML NSsites analysis was carried out with two codon frequency models F3X4 and F61. Analyses were also performed using MEME (Murrell et al., 2012) and FUBAR (Murrell et al., 2013) from Datamonkey (datamonkey.org) (Weaver et al., 2018) to predict rapidly evolving sites. Additional summary of findings is present in the Supplementary file 6.

### Phylogenetic Analysis

MISTR amino acid sequences and related information were retrieved from NCBI, Uniprot, and (Balsa et al., 2012)(Supplementary file 4). Homologs for model species were selected for analysis. Multiple sequence alignment of amino acid sequences were performing using Muscle implemented in Geneious. Phylogenetic analysis was performed using PhyML. Model selection was performed by Smart Model Selection (SMS)(Lefort et al., 2017) integrated into PhyML. The LG +G model was selected for tree building using 1000 bootstrap replicates. FigTree v1.4.2 (http://tree.bio.ed.ac.uk/software/figtree/) was used for tree visualization.

### Cell lines

HeLa, HL-60, L929, Raw 264.7, and HEK293T cell lines were obtained from ATCC. RCC4 (+/-) VHL cell lines were purchased from Sigma. A549 and U2OS cells were generous gifts from Dr. Susan Weiss at the University of Pennsylvania and Dr. Don Gammon at the University of Texas Southwestern Medical Center, respectively. All cell lines except RCC4 were cultured in Corning DMEM with L-Gluatamine, 4.5 g/L Glucose and Sodium Pyruvate supplemented with 10% FBS and 1X Gibco Antibiotic-Antimycotic solution. The Antibiotic-Antimycotic solution was replaced with 0.5 mg/mL G418 in the media for RCC4 cells. All cell lines were maintained at 37°C in a humidified incubator at 5% CO_2_.

### Cell culture treatments

The following were added to cells at the indicated concentrations unless otherwise noted: Staurosporine [1 μM (Abcam)], Interferon Alpha [1000U/mL (PBL Assay Science)], Interferon Gamma [1000U/mL (ThermoFisher)], Actinomycin D [1 μg/mL (Cayman Chemical)], Camptothecin [1 μM (Tocris)], Deferoxamine mesylate [300 μM (Abcam)].

### RT-PCR

Total RNA was extracted using the *Quick*-RNA Miniprep Kit (Zymo) according to the manufacturer’s instructions. 1μg of total RNA was reverse-transcribed using the Maxima First Strand cDNA Synthesis Kit (ThermoFisher) for 10 min at 25°C, 30 min at 50°C, 85°C 5 min. The 20 μL cDNA reaction was subsequently diluted with water to a final volume of 100 μL. 1-2 μL of cDNA was used for 25 μL PCR reactions using the GoTaq Hot Start Master Mix (Promega). Cycling parameters consisted of an initial denaturation of 95°C for 2 min., followed by 28-30 cycles of 95°C for 30s, 50°C for 30s, 72°C for 30s finishing with a final elongation of 72°C 2 min. 20 μL of each PCR product was resolved by 2% agarose gel electrophoresis and visualized using ethidium bromide.

### CRISPR Knockouts

For *MISTRAV* KOs, DNA oligos encoding guide RNAs (gRNA) were synthesized (IDT) and cloned into pSpCas9(BB)-2A-Puro vectors (gift from Feng Zhang, Addgene #62988) according to the protocol here (46). Guide RNAs were positioned in exon 2 (long isoform) with the expectation based on rules of nonsense-mediated decay such that frame-shifts here would be predicted to disrupt the *MISTRAV* ORF while maintaining expression of *miR-147b*. A549 cells were transfected with the gRNA construct, followed by puromycin (Invivogen) selection. Subsequently, limited dilution was performed to establish clonal cell lines. Clones of interest were identified by PCR on genomic DNA harvested with the *Quick*-DNA Miniprep Kit (Zymo) from expanded cell lines using primers flanking exon 2 followed by Sanger sequencing of amplicons by Genewiz. For *MISTR1* KOs, guide RNAs (IDT) were transfected with Cas9 and tracRNA from IDT into A549 cells. Clones were isolated via limiting dilution.

gRNAs were designed using crispr.mit.edu and idt.com

### vMISTRAV stable cell line

vMISTRAV was synthesized (IDT) as a gene block with a C-terminal HA tag and cloned into pMSCV PIG (Puro IRES GFP empty vector) - a gift from David Bartel (Addgene plasmid # 21654). Retroviruses were generated using the retroPack system (Takara) according to manufacturer’s instructions. Following infection of A549 cells, puro selection was performed to select for vMISTRAV-expressing cells.

### Western Blot Analysis

Cells were collected and lysed with RIPA Lysis and Extraction Buffer (ThermoFisher) supplemented with 1X Halt Protease Inhibitor Cocktail (ThermoFisher). For the HIF1a Western blots, nuclear fractions were extracted using Abcam’s Nuclear Fractionation Protocol. Cells cultured in 10-cm dishes were scraped in 500 μL of ice-cold Buffer A (10 mM HEPES, 1.5 mM MgCl2, 10 mM KCl, 0.5 mM DTT, 0.05% NP40, pH 7.9, 1X Halt Protease Inhibitor Cocktail), transferred to 1.5 mL microcentrifuge tubes, and incubated on ice for 10 min. Lysates were centrifuged at 4°C at 3,000 rpm for 10 minutes. Each pellet was resuspended in 374 μL ice-cold Buffer B (5 mM HEPES, 1.5 mM MgCl2, 0.2 mM EDTA, 0.5 mM DTT, 26% glycerol (v/v), pH 7.9, 1X Halt Protease Inhibitor Cocktail) and 26 μL of 4.6M NaCl (final NaCl concentration: 300 mM), homogenized using a syringe with a narrow-gauge needle (27G), and incubated on ice for 30 minutes. Lysates were centrifuged at 4°C at 24,000 x g for 20 min. The supernatant containing the nuclear fraction was transferred to a new tube. Protein concentrations of the extracts were measured using a Bradford assay. Protein samples were subjected to SDS-PAGE and wet-transferred to a 0.2 μM Immobilon-PSQ PVDF membrane (Millipore) at 200 mA for 90 minutes. Membranes were blocked with blocking buffer (5% BSA or milk in TBST) for 1 hour at RT, and then incubated with primary antibodies at 4°C overnight. The following primary antibodies were used: SDHA (D6J9M) XP Rabbit mAB (CST), PARP (CST), NDUFA4 (ThermoFisher), IDO (Novus Biologicals), C15orf48 (Aviva Systems Biology), HA (Sigma), NDUFA4L2 (ThermoFisher), HIF1α (Proteintech), HDAC1 (Proteintech), β-actin (Sigma), and beta-3 Tubulin (ThermoFisher). Membranes were washed three times with TBST and then incubated with secondary antibodies for 1 hour at RT. Goat Anti-Rabbit IgG (Bio-Rad) and Goat Anti-Mouse IgG (Bio-Rad) were used as secondary antibodies. Membranes were washed three times with TBST and then incubated with Pierce ECL Plus Western Blotting Substrate (ThermoFisher). Blots were imaged using the ChemiDoc MP Imager (Bio-Rad).

### miRNA qPCR

Total RNA was extracted from cultured cells using the mirVana miRNA Isolation kit (Ambion) following the manufacturer’s protocol. For each sample, 10 ng of total RNA was used as input for cDNA synthesis using the TaqMan Advanced miRNA cDNA Synthesis Kit (Thermofisher). *hsa-miR-147b-3p* and *hsa-miR-210-3p* levels were assessed by TaqMan Advanced miRNA Assays (Thermofisher) and TaqMan Fast Advanced miRNA master mix (Thermofisher). *hsa-mir-423-5p* (Thermofisher) served as endogenous control for analysis of miRNA expression. PCR was run in an Applied Biosytems QuantStudio 7 Real-Time PCR instrument following the manufacturer’s instructions.

### Cell-viability assays

A549 cells were plated at 1 x 10^4^ cells/well in opaque white 96-well plates (Corning) in 100 μL of media. 24 hours later, spent medium was aspirated and replaced with 75 μL of fresh media supplemented with 1000 U/mL IFN-γ (ThermoFisher). 24 hours following IFN-γ addition, 25 μL of media containing STS (Abcam) was added (final STS treatment concentration: 1 μM). 16 hours later, cell viability was assessed using CellTiter-Glo (Promega) following the manufacturer’s instructions.

### IncuCyte analysis of Caspase 3/7 activity

For experiments on the *MISTRAV* KO clones, 5 x 10^3^ cells were seeded and primed with IFN-γ as above. 24 hours post IFN-γ addition, 25 μL of media containing STS and CellEvent Caspase-3/7 Green Detection Reagent (ThermoFisher) at final treatment concentrations of 1 μM and 2.5 μM, respectively, was added. For experiments on the *MISTR1* KO and the vMISTRAV cell lines, 5 x 10^3^ cells/well were plated in opaque white 96-well plates (Corning) in 75 μL of media. 24 hours later, 25 μL of media containing the appropriate drug and Caspase-3/7 detection reagent was added (*MISTR1* KO cell lines: 1 μM STS, 2.5 μM CellEvent Caspase-3/7 Green Detection Reagent; EV and vMISTRAV cell lines: 1 μM STS, 1 μg/mL ActD, 1 μM CPT, 5 μM IncuCyte Caspase-3/7 Red Apoptosis Assay Reagent). To determine the cell number at the initial treatment timepoint, 25 μL of media containing Vybrant DyeCycle Green Stain or SYTO 60 Red Fluorescent Nucleic Acid Stain (final concentration: 1 μM) was added to a set of wells for each cell line. Plates were placed in an IncuCyte S3 Live-Cell Analysis System (Essen Bioscience) with a 10X objective in a standard cell culture incubator at 37°C and 5% CO_2_. Four images/well were collected every 2 hours in phase-contrast and fluorescence. The integrated object counting algorithm was used to count fluorescent objects/mm^2^ for each time point. Percent apoptosis was determined by dividing the number of caspase-3/7 objects/mm^2^ at each time point by the number of cells/mm^2^ at the initial treatment timepoint.

### Chemical Hypoxia Induction

A day after plating cells in 6-well plates or 10-cm dishes, chemical hypoxia was induced by treating cells with 300 μM DFO. 24 hours later cells were either collected in RIPA buffer or subjected to nuclear fractionation protocol as described above.

### miRNA and MRE analysis

Predicted MREs in MISTR1 were retrieved from Targetscan (Agarwal et al., 2015) and mirDB (Wong and Wang, 2015). miRNA and MISTR1 sequences were retrieved from NCBI (Supplementary file 6).

### Transfection of miRNAs and miRNA reporter luciferase assays

293T cells were seeded at 1 x 10^4^ cells/well in opaque white 96-well plates (Corning) in 75μL of media. The next day, cells were transfected with 50 ng/well of the psiCHECK-2 (Promega) construct using the FuGENE HD Transfection Reagent (Promega), following the manufacturer’s instructions. 24 hours later, cells were transfected with 1 pmol/well of miRNA mimics (ThermoFisher) using Lipofectamine RNAiMAX Transfection Reagent (ThermoFisher) according to manufacturer’s instructions. The following miRNA mimics were used: *hsa-miR-210-3p*, *hsa-miR-7-5p*, *hsa-miR-202-5p*, *hsa-miR-145-5p*, *hsa-miR-205-5p*, *hsa-miR-147b-3p*, and Negative Control #1 (ThermoFisher). 48 hours after miRNA transfection, firefly and *Renilla* luciferase activities were measured using the Dual-glo Luciferase assay (Promega).

### Constructs

hMISTRAV-GFP and vMISTRP-GFP vectors were generated via PCR cloning. Briefly, hMISTRAV and vMISTRAV were synthesized as gBlocks (IDT), amplified using primers with KpnI and BamHI RE sites with Phusion Master Mix (NEB), digested, and ligated to N1-EGFP (Clontech) digested with KpnI and BamHI. Clones were confirmed by Sanger sequencing. Primer sequences available in Supplementary file 5.

#### Confocal Images

One day following transfection of either 1 μg hMISTRAV-GFP or 1 μg vMISTRAV-GFP, HeLa cells were fixed and stained with mitoTracker Red (Thermo). Images were taken in the confocal microscopy core at the University of Utah.

### Protein Modeling

A recently published predicted structure of Complex IV (PDB:5Z62)(Zong et al., 2018), which contains MISTR1, was used for modeling. The structures of MISTR paralogs (MISTRAV, MISTRH) were predicted using Swiss-Model (Waterhouse et al., 2018). UCSF Chimera (https://www.cgl.ucsf.edu/chimera/)(Pettersen et al., 2004) was used for visualization, mapping rapidly evolving sites, and analysis.

### Statistical analysis

Experimental data are presented at means ± SD. Statistical significance was determined by two-tailed unpaired student’s t-test. GraphPad Prism software (Version 8.3.0) was used for statistical analysis.

## ACKNOWLEDGEMENTS

We thank Malory Monson and Diane Downhour for technical assistance. We express gratitude to other members of the Hancks and Elde Labs as well as Don Gammon for feedback and discussion through the course of this project. We also thank Dan Propheter and Mike O’Donnell for comments on the manuscript.

## ADDITIONAL INFORMATION

### Funding

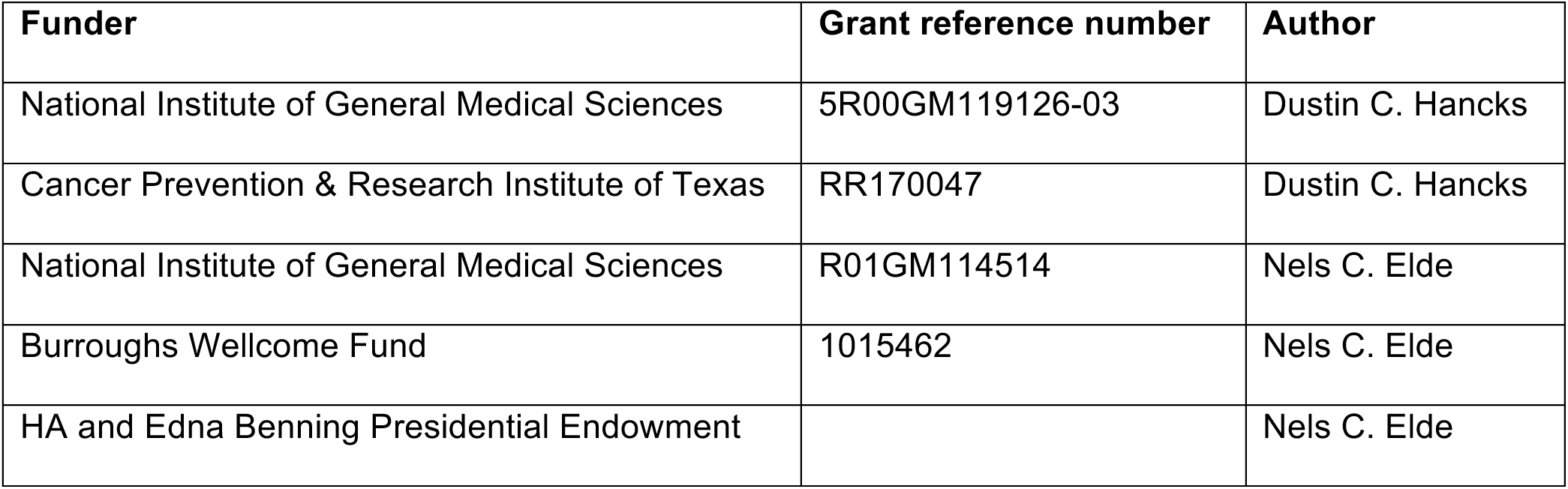

### Author Contributions

Conceptualization, M.S., N.C.E, and D.C.H.; Methodology, M.S., N.C.E, and D.C.H.; Validation, M.S. and D.C.H.; Investigation, M.S., T.C., C.P., P.J., and D.C.H.; Resources, M.S., T.C., C.P., P.J., N.C.E. and D.C.H.; Writing – Original Draft, M.S. and D.C.H.; Writing – Review & Editing, M.S., N.C.E., and D.C.H.; Visualization, M.S. and D.C.H.; Supervision, N.C.E. and D.C.H.; Funding Acquisition, N.C.E. and D.C.H.

## ADDITIONAL FILES

### Supplementary files

Supplementary file 1. Nucleotide sequence information for rapid evolution analysis

Supplementary file 2. *miR-147b* target prediction output from miRDB

Supplementary file 3. *miR-147b* target prediction output from TargetScan

Supplementary file 4. Sequence information for evolutionary analysis of MISTR homologs

Supplementary file 5. Primers and oligos used in this study

Supplementary file 6. Supplementary file 6

Supplementary file 7. Key Resources Table

---

**Figure 1- figure supplement 1:**
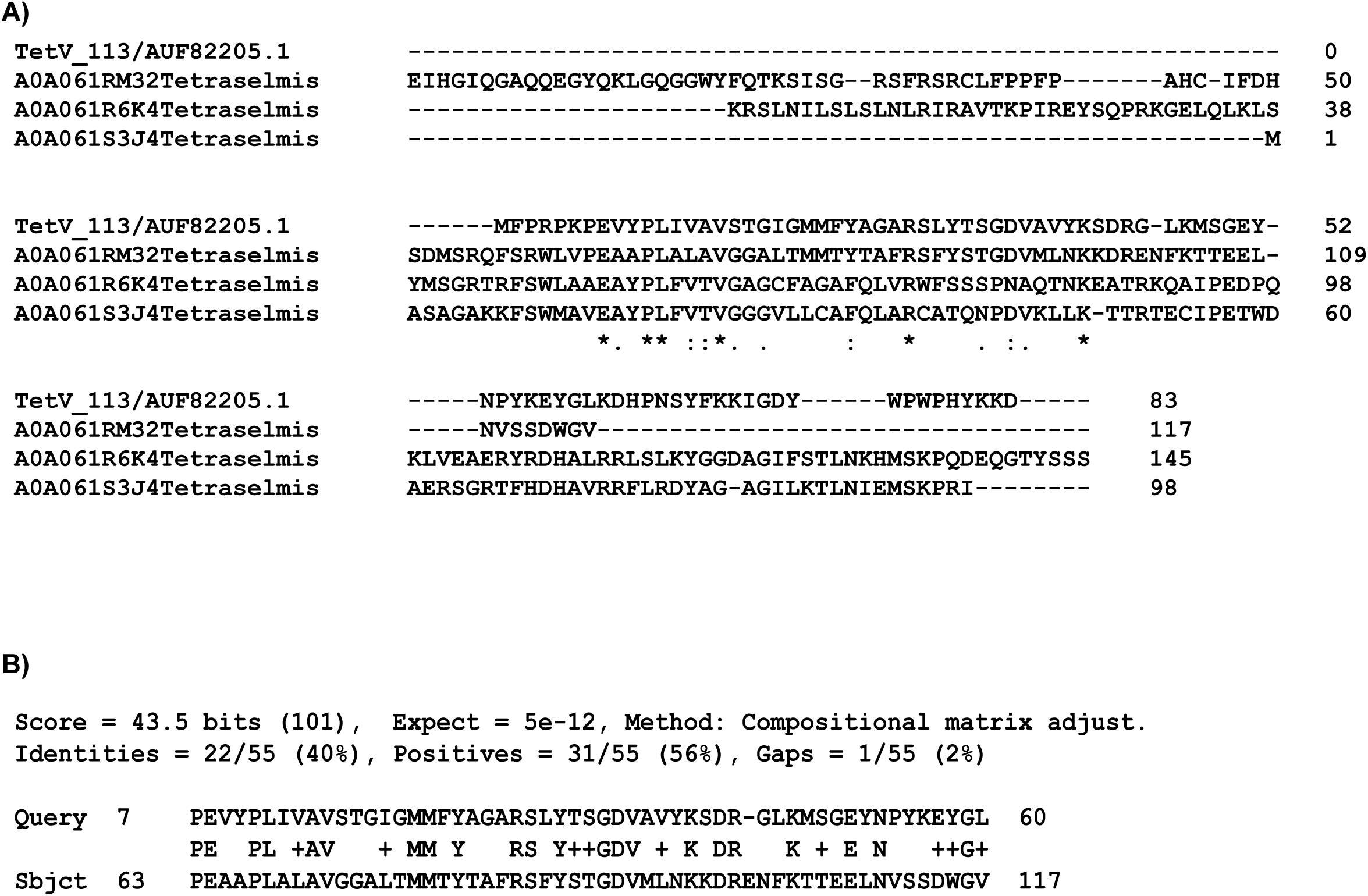
Sequence analysis of Tetraselmis virus 1 MISTR (vMISTRA). **A)** Clustal omega amino acid alignment of Tetraselmis virus 1 MISTR with three Tetraselmis MISTR protein sequences from the database. **B)** blastp analysis of Tetraselmis virus 1 MISTR - Query - with Tetraselmis MISTR (A0A061RM32) - Subject.

**Figure 2- figure supplement 1:**
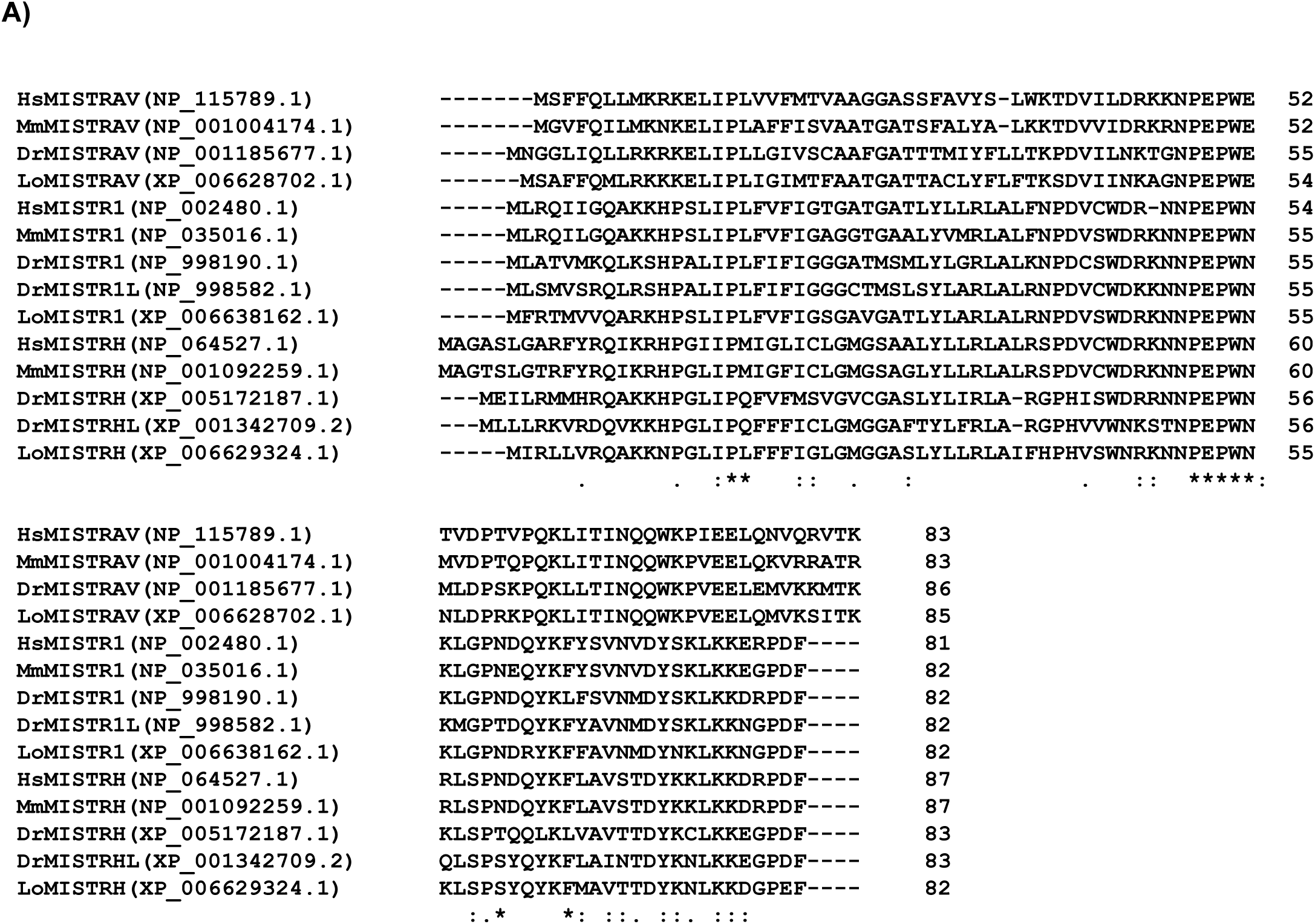
MISTR factors are conserved in vertebrates. Clustal omega amino acid alignment of MISTRAV, MISTR1, and MISTRH sequences. Hs - *Homo sapiens* (Human), Mm - *Mus musculus* (mouse), Dr - *Danio rerio* (zebrafish), Lo - *Lepisosteus oculatus* (spotted-gar). Accession numbers are for NCBI.

**Figure 3- figure supplement 1:**
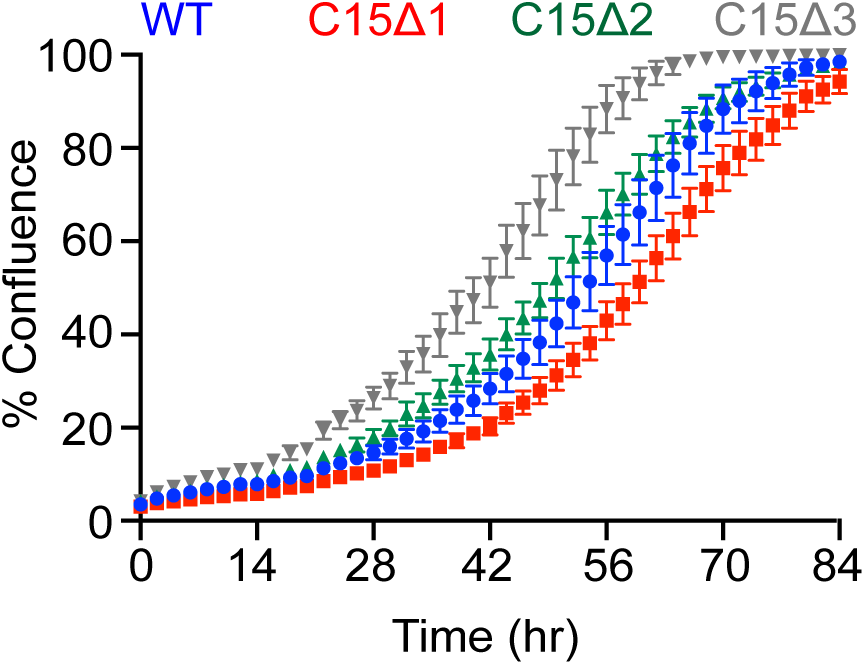
Proliferation rates of A549 C15 (*MISTRAV* KO) knockout clonal lines measured using IncuCyte. Changes in % confluence were used as a surrogate marker of cell proliferation. Data represent means ± SD (n=6 replicates).

**Figure 4- figure supplement 1:**
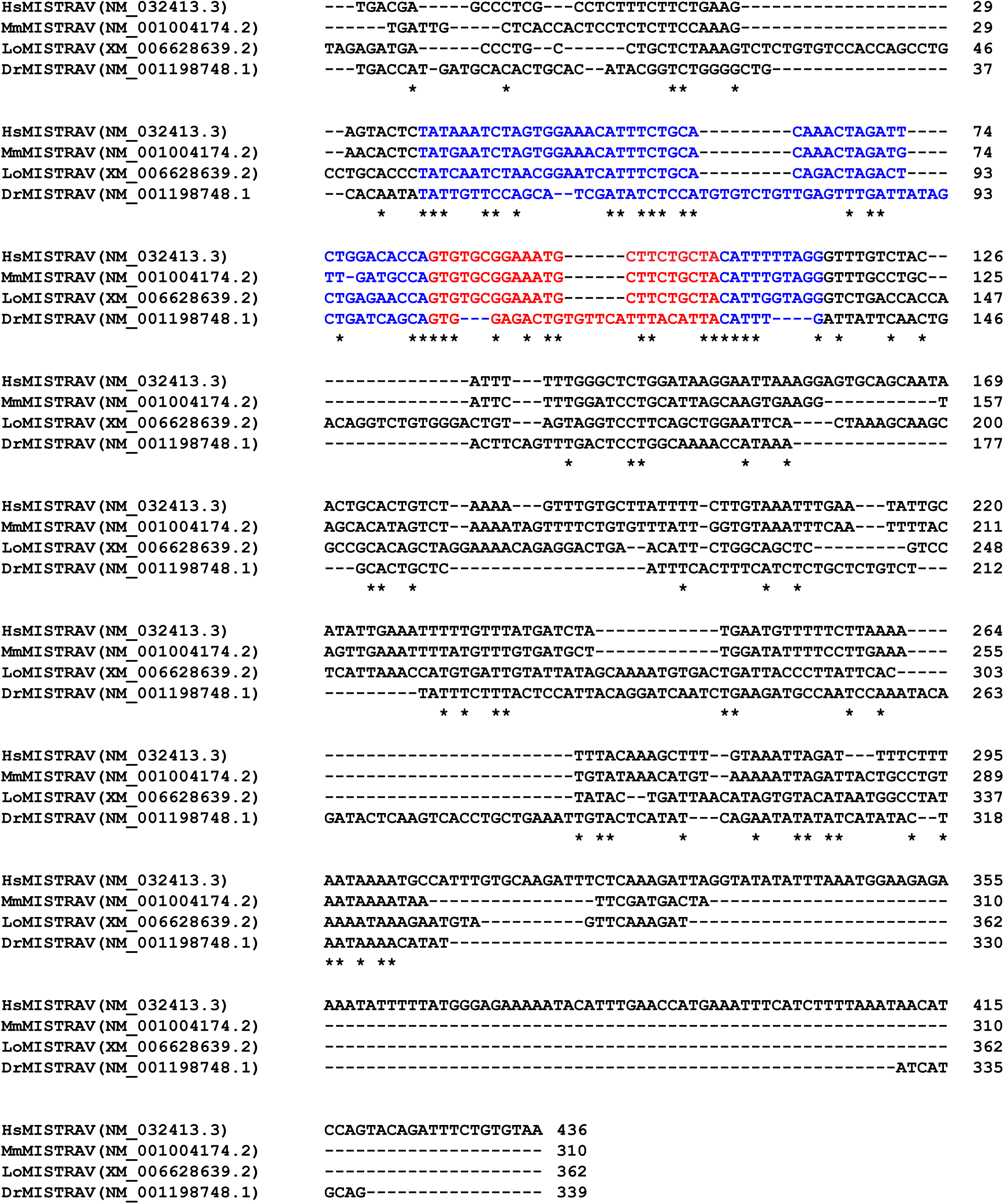
Zebrafish lack intact *miR-147b*. Clustal omega nucleotide alignment of *MISTRAV* 3’-UTR sequences. Alignment starts with MISTRAV stop codon. Predicted *pre-mir-147b* (blue) relative to human annotation, predicted *miR-147b* (red). Hs - *Homo sapiens* (Human), Mm - *Mus musculus* (mouse), Dr - *Danio rerio* (zebrafish), Lo - *Lepisosteus oculatus* (spotted-gar). Accession numbers are for NCBI.

**Figure 5- figure supplement 1:**
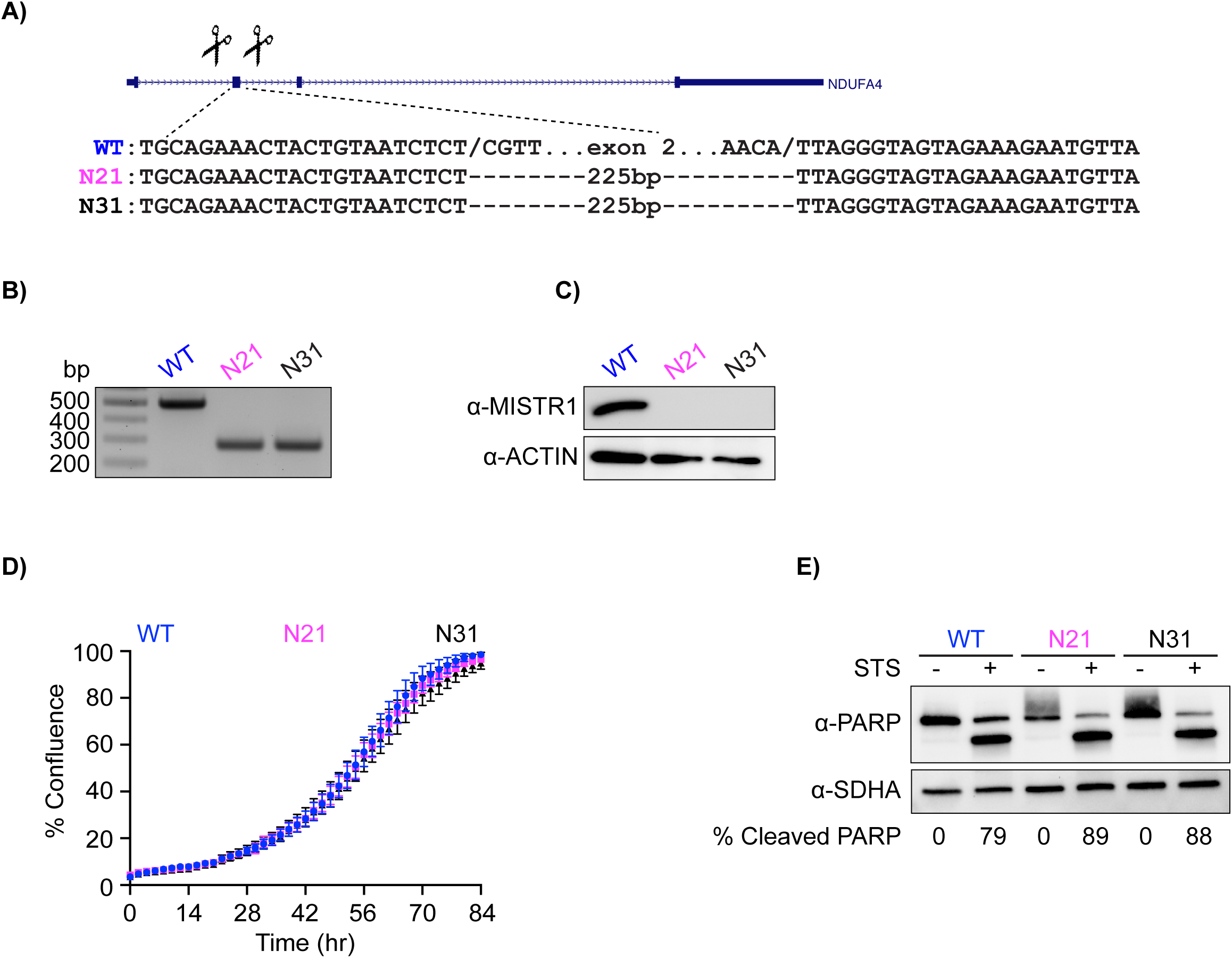
Generation and characterization of *MISTR1* KO A549 cells. **A)** CRISPR/CAS deletion strategy for *MISTR1*. Scissors indicate relative locations of guide RNAs designed to target sequences flanking exon 2 of this gene. The exon 2 deletion strategy was employed for ease of genotyping. Gene structure from UCSC genome browser. Sequences of breakpoints identified a 225bp deletion that included exon 2. Note identical repaired breakpoints were recovered for both clones. **B)** Agarose gel resolving amplicons from genotyping PCR of A549 KO clones. **C)** Western blot analysis using lysates from WT and *MISTR1* KO clones. **D)** Measurement of proliferation rates using IncuCyte for *MISTR1* KO A549 cell line. Changes in % confluence were used as a surrogate marker of cell proliferation. Data represent means ± SD (n=6 replicates). **E)** Western blot analysis of cleaved PARP levels using lysates from WT and *MISTR1* KO cells following 16 hours of STS treatment. Densitometry analysis of PARP levels was performed using Image Lab version 6.0.1 (Bio-Rad). % Cleaved PARP= (cleaved PARP/(Full + Cleaved PARP)) * 100. STS - staurosporine.

**Figure 7- figure supplement 1:**
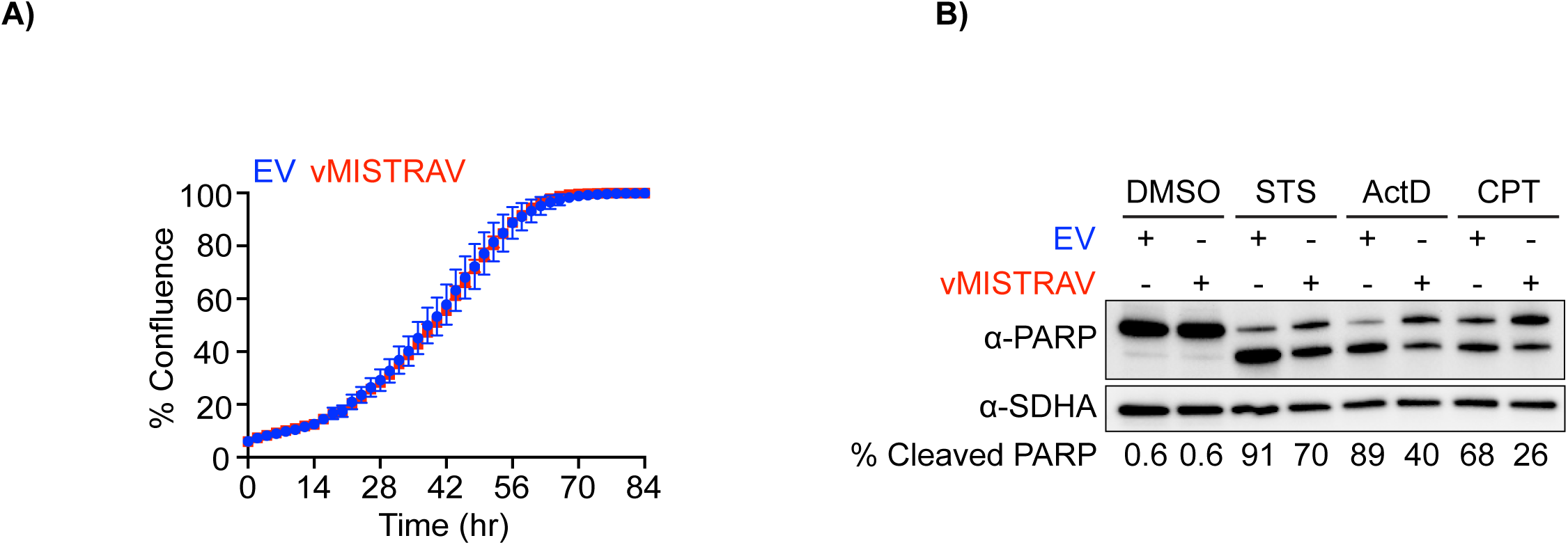
Characterization of WT A549 cells stably-expressing vMISTRAV. **A)** Proliferation rates of empty vector (EV) and vMISTRAV-expressing cells measured using IncuCyte. Changes in % confluence were used as a surrogate marker of cell proliferation. Data represent means ± SD (n=6 replicates). **B)** Western blot analysis of cleaved PARP levels using lysates from EV and vMISTRAV-expressing cells following treatment with activators of apoptosis. STS - staurosporine, ActD - actinomycin D, CPT - camptothecin. Lysates were collected 16 hours after treatment with STS or ActD and 24 hours after treatment with CPT. Densitometry analysis of PARP levels was performed using Image Labversion 6.0.1 (Bio-Rad). % Cleaved PARP= (cleaved PARP/(Full + Cleaved PARP)) * 100.

